# The Potential for SARS-CoV-2 to Evade Both Natural and Vaccine-induced Immunity

**DOI:** 10.1101/2020.12.13.422567

**Authors:** Emily Shang, Paul H. Axelsen

**Author notes:** To whom correspondence should be addressed: Paul H. Axelsen, Department of Pharmacology, 1009C Stellar Chance Laboratories, University of Pennsylvania, Philadelphia PA, 19104-6084.

## Abstract

SARS-CoV-2 attaches to the surface of susceptible cells through extensive interactions between the receptor binding domain (RBD) of its spike protein and angiotensin converting enzyme type 2 (ACE2) anchored in cell membranes. To investigate whether naturally occurring mutations in the spike protein are able to prevent antibody binding, yet while maintaining the ability to bind ACE2 and viral infectivity, mutations in the spike protein identified in cases of human infection were mapped to the crystallographically-determined interfaces between the spike protein and ACE2 (PDB entry 6M0J), antibody CC12.1 (PDB entry 6XC2), and antibody P2B-2F6 (PDB entry 7BWJ).

Both antibody binding interfaces partially overlap with the ACE2 binding interface. Among 16 mutations that map to the RBD:CC12.1 interface, 11 are likely to disrupt CC12.1 binding but not ACE2 binding. Among 12 mutations that map to the RBD:P2B-2F6 interface, 8 are likely to disrupt P2B-2F6 binding but not ACE2 binding. As expected, none of the mutations observed to date appear likely to disrupt the RBD:ACE2 interface.

We conclude that SARS-CoV-2 with mutated forms of the spike protein may retain the ability to bind ACE2 while evading recognition by antibodies that arise in response to the original wild-type form of the spike protein. It seems likely that immune evasion will be possible regardless of whether the spike protein was encountered in the form of infectious virus, or as the immunogen in a vaccine. Therefore, it also seems likely that reinfection with a variant strain of SARS-CoV-2 may occur among people who recover from Covid-19, and that vaccines with the ability to generate antibodies against multiple variant forms of the spike protein will be necessary to protect against variant forms of SARS-CoV-2 that are already circulating in the human population.

## Introduction

The receptor binding domain (RBD) of the human coronavirus SARS-CoV-2 spike (S) protein binds to angiotensin converting enzyme 2 (ACE2) on the surface of susceptible cells. A crystal structure of the RBD:ACE2 complex shows that the binding interface involves 37 amino acid residues and a surface area of 1687 Å^2^, at the end of the spike protein opposite from its membrane anchor (figure 1a).^1^ As expected for a protein that has a critical role in cell attachment, the RBD sequence is highly conserved, making the RBD an attractive target for vaccine development.^2-5^ Sequences for over 100,000 SARS-CoV-2 isolates from around the world are available in the GISAID^6^ database, and mutations in the spike protein may be identified using COVID CG interface.^7^ It is of interest to map the locations of these mutations on the spike protein surface because mutations in the RBD:ACE2 interface indicate sites where at least some mutations are tolerated, while mutations elsewhere may interfere with antibody binding.

**Figure 1:**
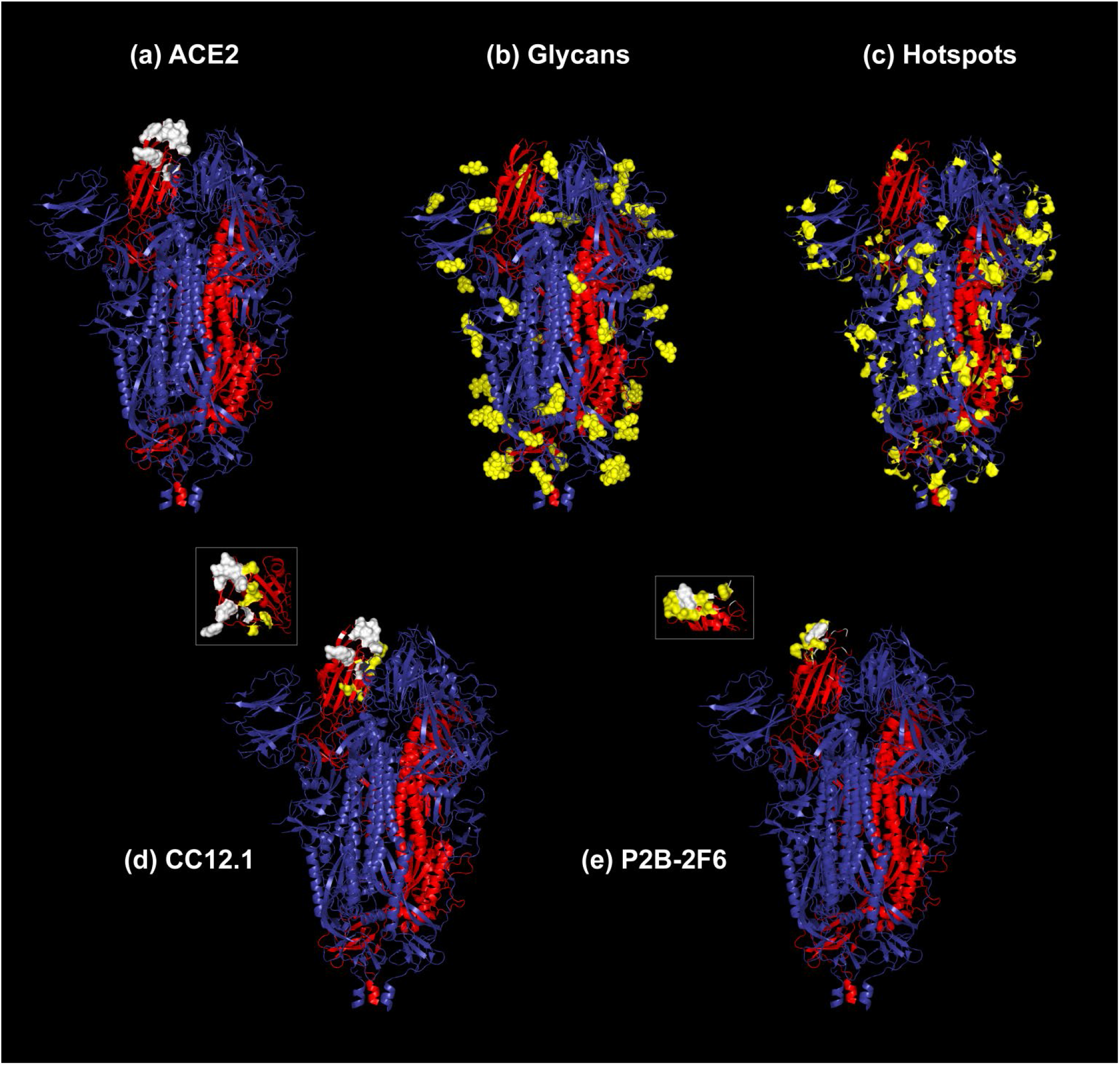
Surface features of the trimeric SARS-CoV-2 spike protein. In each panel, one of the trimers is red, while the other two are blue. (a) The RBD surface that interfaces with the ACE2 receptor is rendered in white. Only one of the three interfaces in the trimer has been rendered. (b) Glycosylation sites are rendered in yellow. (c) Mutated residues identified in circulating viruses are rendered in yellow. (d) The RBD surface that interfaces with the CC12.1 antibody is rendered in yellow, and with both the CC12.1 antibody and ACE2 is rendered in white. Only one of the three interfaces in the trimer has been rendered. *Inset*: alternative view of the RBD surface. (e) The RBD surface that interfaces with the P2B-2F6 antibody is rendered in yellow, and with both the P2B-2F6 antibody and ACE2 is rendered in white. Only one of the three interfaces in the trimer has been rendered. *Inset*: alternative view of the RBD surface.

Natural infection with SARS-CoV-2 generates multiple neutralizing antibodies against the RBD, including those designated H014,^8^ P2B-2F6,^9^ CC12.1,^10^ CC12.3,^10^ COVA2-15,^11^ CB6,^12^ and B38.^13^ X-ray crystal structures are available for P2B-2F6, CC12.1, CC12.3, B38, and CB6 in complex with the RBD. The binding interfaces for CC12.1, CC12.3, B38, and CB6 largely overlap with each other, and with the binding interface for ACE2.^14-16^ Only CC12.1 was examined in this work because its binding interface was largest (figure 1d). We also examined the interface between the RBD and P2B-2F6 because it differs significantly from that of CC12.1 (figure 13).^15^

We hypothesized that mutations would either not map to the RBD:ACE2 interface, or they would represent conservative substitutions unlikely to disrupt the RBD:ACE2 interface. The aim of the present study was to test this hypothesis and determine whether mutations already circulating in the human population mapped to the RBD:CC12.1 and RBD:P2B-2F6 interfaces and were likely to disrupt the binding of those antibodies to the RBD.

## Methods

Mutations in the SARS-Cov-2 spike protein sequence were identified using the COVID CG interface^7^ to the GISAID database^6^ which, at the time of this study, contained 147,045 sequences. Protein binding interfaces were defined by calculating the accessible surface area (SASA) of the unmutated residue with VMD^17^ and a 1.4Å probe. Residues were designated to be “in” the binding interface if they had a SASA when the proteins were apart that became zero when the proteins were considered together. Residues were designated to be on the “edge” of an interface if they had a SASA when the proteins were apart that became smaller when the proteins were considered together. Mutation sites were examined and figures were generated using PyMol.^18^ Mutations were designated as “disruptive” if they increased sidechain volume or changed electrostatic character, “destabilizing” if they created hydrogen bond donors or acceptors that could not be satisfied by the mutation or solvent accessibility, and “permissive” if they did not increase side chain volume, change electrostatic character, or create unsatisfied hydrogen bond donors or acceptors. Figures illustrating the entire ectodomain of the spike protein are based on the cryo-EM structure (PDB entry 6VXX).^19^

## Results

We identified 12 mutations in the RBD:CC12.1 interface (table 1) and 12 mutations on the edge of this interface (table 2). We also identified 11 mutations in the RBD:P2B-2F6 interface (table 3), and 9 mutations on the edge of this interface (table 4). The protein-protein interfaces was examined at the site of each mutation to determine whether the observed mutation altered hydrogen bonding or electrostatic pairs, and whether the interface could accommodate the volume and shape changes implied by the mutation.

**Table 1:**
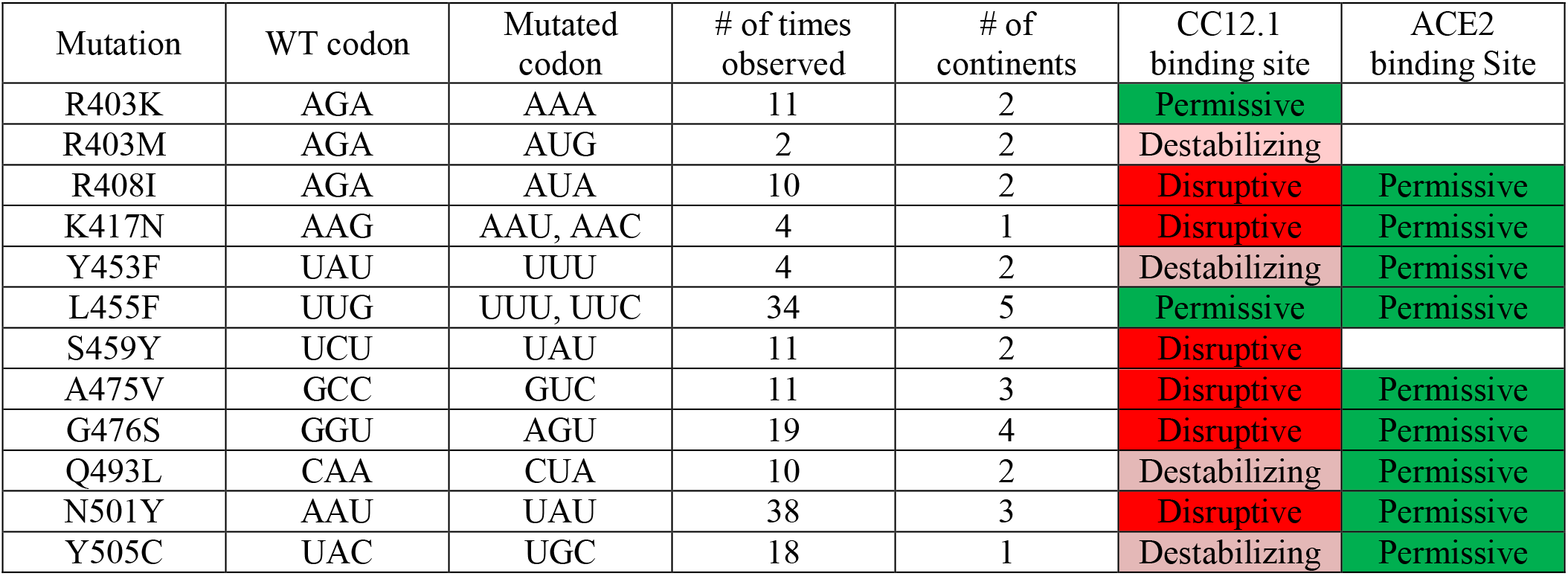
Mutations in the RBD:CC12.1 Interface

| Mutation | WT codon | Mutated codon | # of times observed | # of continents | CC12.1 binding site | ACE2 binding Site |
| --- | --- | --- | --- | --- | --- | --- |
| R403K | AGA | AAA | 11 | 2 | Permissive |  |
| R403M | AGA | AUG | 2 | 2 | Destabilizing |  |
| R408I | AGA | AUA | 10 | 2 | Disruptive | Permissive |
| K417N | AAG | AAU, AAC | 4 | 1 | Disruptive | Permissive |
| Y453F | UAU | UUU | 4 | 2 | Destabilizing | Permissive |
| L455F | UUG | UUU, UUC | 34 | 5 | Permissive | Permissive |
| S459Y | UCU | UAU | 11 | 2 | Disruptive |  |
| A475V | GCC | GUC | 11 | 3 | Disruptive | Permissive |
| G476S | GGU | AGU | 19 | 4 | Disruptive | Permissive |
| Q493L | CAA | CUA | 10 | 2 | Destabilizing | Permissive |
| N501Y | AAU | UAU | 38 | 3 | Disruptive | Permissive |
| Y505C | UAC | UGC | 18 | 1 | Destabilizing | Permissive |

**Table 2:** Mutations on the edge of the RBD:CC12.1 Interface

| Mutation | WT codon | Mutated codon | # of times observed | # of continents | CC12.1 binding site | ACE2 binding site |
| --- | --- | --- | --- | --- | --- | --- |
| S477N | AGC | AAC | 7166 | 3 | Permissive | Permissive |
| S477G | AGC | GGC | 2 | 1 | Permissive | Permissive |
| S477R | AGC | AGA, AGG | 20 | 3 | Permissive | Permissive |
| S477I | AGC | AUC | 14 | 1 | Permissive | Permissive |
| E484Q | GAA | CAA | 26 | 3 | Permissive | Permissive |
| E484K | GAA | AAA | 26 | 4 | Permissive | Permissive |
| E484A | GAA | GCA | 7 | 2 | Permissive | Permissive |
| F490S | UUU | UCU | 10 | 2 | Permissive | Permissive |
| F490L | UUU | UUA, UUG | 6 | 3 | Permissive | Permissive |
| S494P | UCA | CCA | 80 | 5 | Permissive | Permissive |
| S494L | UCA | UUA | 6 | 2 | Permissive | Permissive |
| V503F | GUU | UUU | 4 | 2 | Permissive | Permissive |

**Table 3:** Mutations in the RBD:P2B-2F6 Interface

| Mutation | WT codon | Mutated codon | # of times Observed | # of continents | P2B-2F6 binding site | ACE2 binding site |
| --- | --- | --- | --- | --- | --- | --- |
| G446V | GGU | GUU | 17 | 4 | Disruptive | Permissive |
| G446S | GGU | AGU | 2 | 1 | Disruptive | Permissive |
| L452M | CUG | AUG | 1 | 1 | Permissive |  |
| V483F | GUU | UUU | 17 | 3 | Disruptive |  |
| V483A | GUU | GCU | 48 | 3 | Permissive |  |
| V483I | GUU | AUU | 3 | 1 | Permissive |  |
| E484Q | GAA | CAA | 26 | 3 | Destabilizing | Permissive |
| E484K | GAA | AAA | 26 | 4 | Destabilizing | Permissive |
| E484A | GAA | GCA | 7 | 2 | Destabilizing | Permissive |
| F490S | UUU | UCU | 10 | 2 | Destabilizing |  |
| F490L | UUU | UUA, UUG | 6 | 3 | Permissive |  |

**Table 4:** Mutations on the edge of the RBD:P2B-2F6 Interface

| Mutation | WT codon | Mutated codon | # of times observed | # of continents | P2B-2F6 binding site | ACE2 binding site |
| --- | --- | --- | --- | --- | --- | --- |
| R346K | AGA | AAA | 4 | 3 | Permissive |  |
| K444R | AAG | AGG | 18 | 1 | Permissive |  |
| K444N | AAG | AAU, AAC | 6 | 3 | Permissive |  |
| V445A | GUU | GCU | 4 | 4 | Permissive | Permissive |
| G482S | GGU | AGU | 4 | 2 | Permissive |  |
| G485R | GGU | Varies | 33 | 1 | Permissive | Permissive |
| Q493L | CAA | CUA | 10 | 2 | Permissive | Permissive |
| S494P | UCA | CCA | 80 | 5 | Permissive | Permissive |
| S494L | UCA | UUA | 6 | 2 | Permissive | Permissive |

### Mutations in the RBD:CC12.1 interface (table 1)

R403 is in the interface with CC12.1, but it does not interact directly with any ACE2 residues (Fig 2b). In the interface with CC12.1, R403-Nη2 forms a hydrogen bond with N92-O in the CC12.1 light chain (Fig 2a). R403K is a conservative mutation with respect to electrostatic properties, and its sidechain volume is slightly smaller. R403M is nonconservative, and is the result of a double mutation. It appears that both R403K and R403M are may be tolerated in the interface with ACE2 (i.e. the mutations are permissive with respect to this interaction). R403K can maintain the hydrogen bond with N92-O in the interface with CC12.1 and may be permissive in that interface, but R403M is likely to be destabilizing.

**Figure 2:**
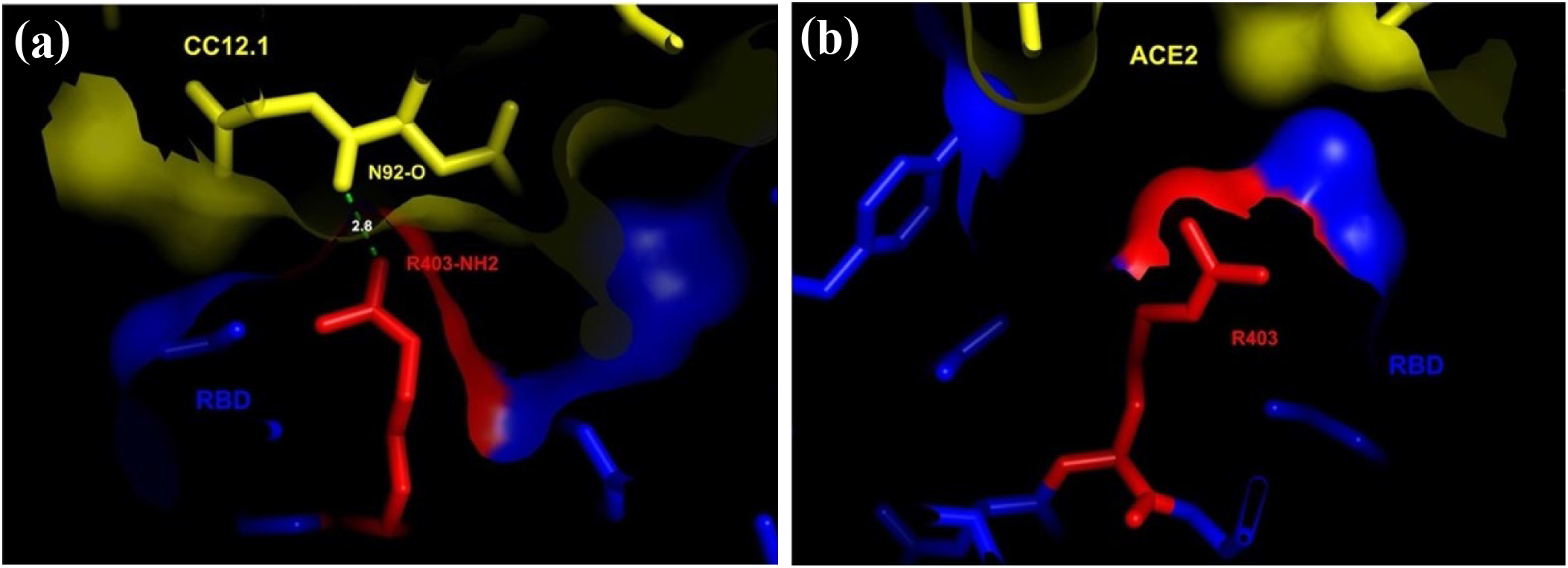
R403K/M. (a) RBD:CC12.1 interface. (b) RBD:ACE2 interface.

R408 is in the interface with CC12.1, but not the interface with ACE2 (Fig 3). The sidechain of R408 forms a hydrogen bond with the sidechain of Y94 (Fig 3a). R408-Nη2 forms a hydrogen bond with Y94-Oη of the CC12.1 light chain. Therefore, R408I is nonconservative and likely to be disruptive in the interface with CC12.1.

**Figure 3:**
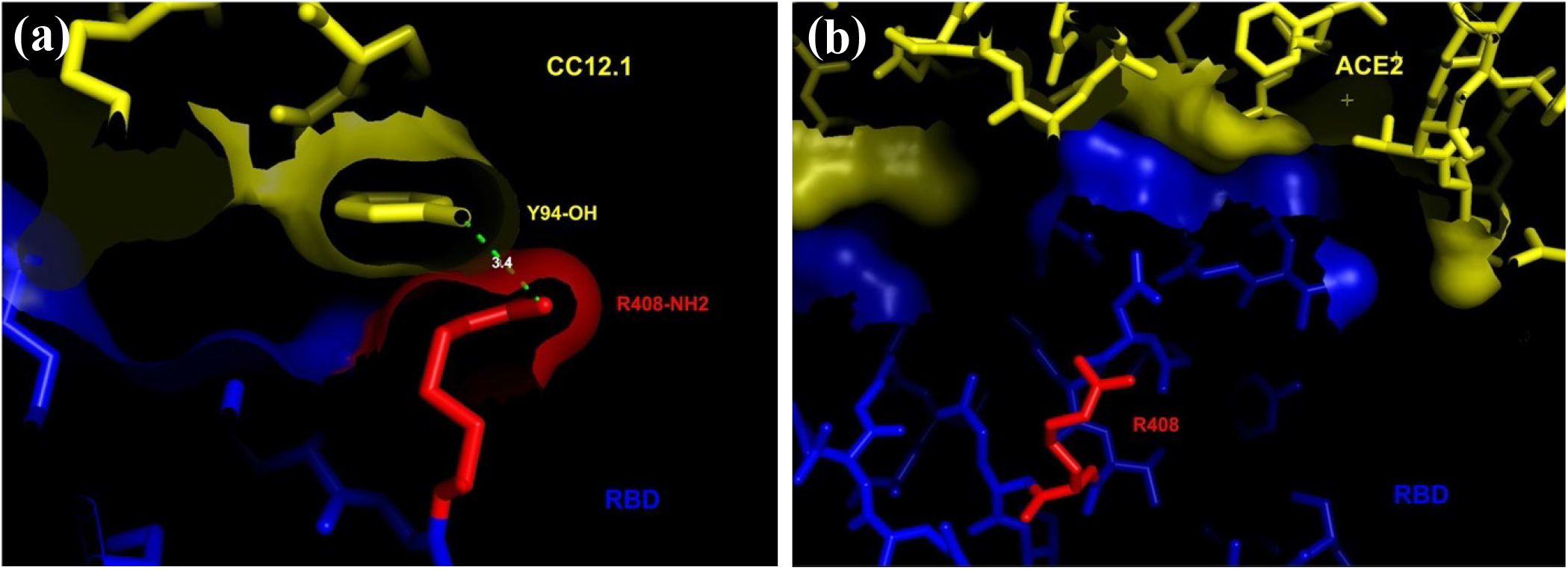
R408I. (a) RBD:CC12.1 interface. (b) RBD:ACE2 interface.

K417 is in the interface with CC12.1, and on the edge of the interface with ACE2 (Fig 4). K417-Nζ forms a hydrogen bond with the carboxylate group of D97 on the CC12.1 heavy chain (Fig 4a), and with the carboxylate group of D30 in ACE2 (Fig 4b). K417N is nonconservative with respect to electrostatic charge, shape, and size of the side chain. Therefore, it is likely to be disruptive in the interface with CC12.1, but permissive on the edge of ACE2 interface.

**Figure 4:**
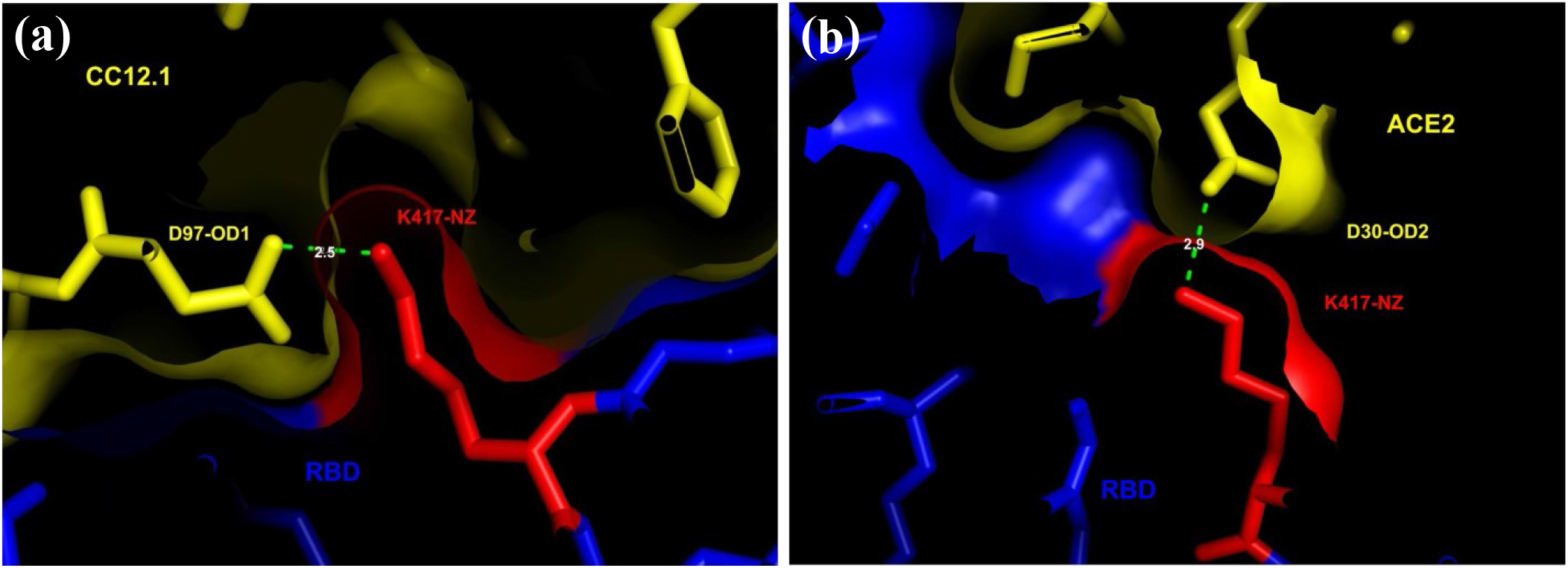
K417N. (a) RBD:CC12.1 interface. (b) RBD:ACE2 interface.

Y453 is in the interfaces with CC12.1 and ACE2 (Fig 5). In the RBD:CC12.1 interface, Y453-Oη appears to hydrogen bond with N92-Nδ2 in the CC12.1 heavy chain (Fig 5a), and with the aromatic ring face of H34 in ACE2 (Fig 5b). Y453F is generally considered a conservative mutation, however the loss of a hydrogen bonds in the interface with CC12.1 is likely to be destabilizing, while it may still form an edge-face interaction with H34 in the interface with ACE2.

**Figure 5:**
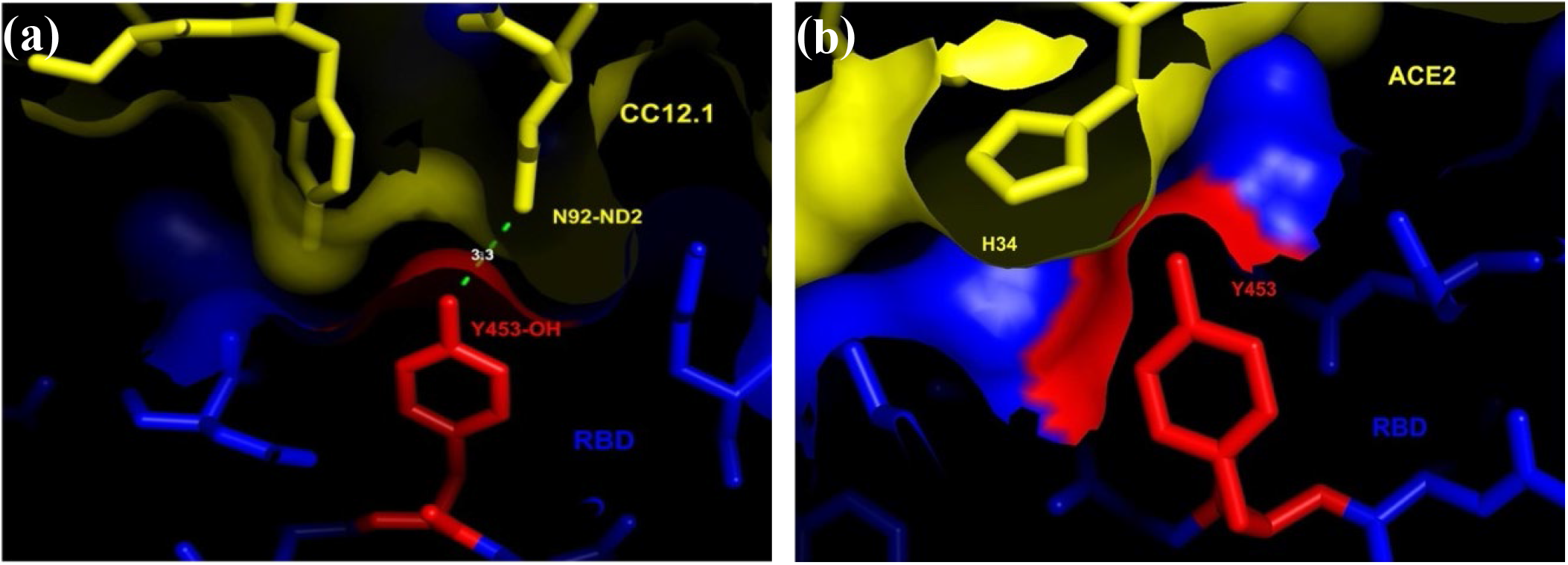
Y453F. (a) RBD:CC12.1 interface. (b) RBD:ACE2 interface.

L455 is in the interfaces with CC12.1 and ACE2 (Fig 6). In the RBD:CC12.1 interface, L455-O forms a hydrogen bond with Y33-Hη of the CC12.1 heavy chain, while the aliphatic side chain of L455 makes extensive VdW contact with the aliphatic side chain of D97 (Fig 6a). In the RBD:ACE2 interface, the aliphatic side chain of L455 makes VdW contact with the side chain of H34 (Fig 6b). L455F entails a different shape and slightly larger volume, but there appears to be sufficient room for these chains in both interfaces for this mutation to be permissive.

**Figure 6:**
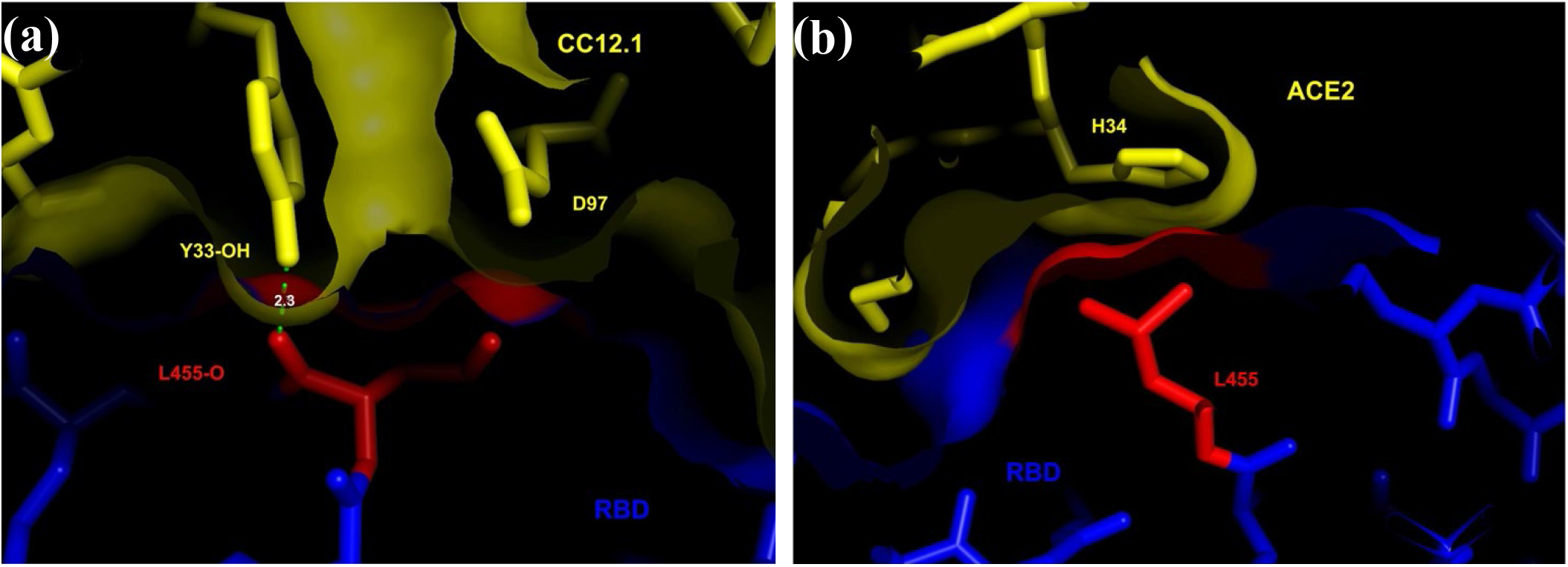
L455F. (a) RBD:CC12.1 interface. (b) RBD:ACE2 interface.

S459 is in the interface with CC12.1, but not the interface with ACE2 (Fig 7). In the RBD:CC12.1 interface, the main chain of S459 makes VdW contact with the main chain of G54 in the CC12.1 heavy chain (Fig 7a). S459Y is likely to disrupt the interface with CC12.1, as there is little room for the much larger side chain.

**Figure 7:**
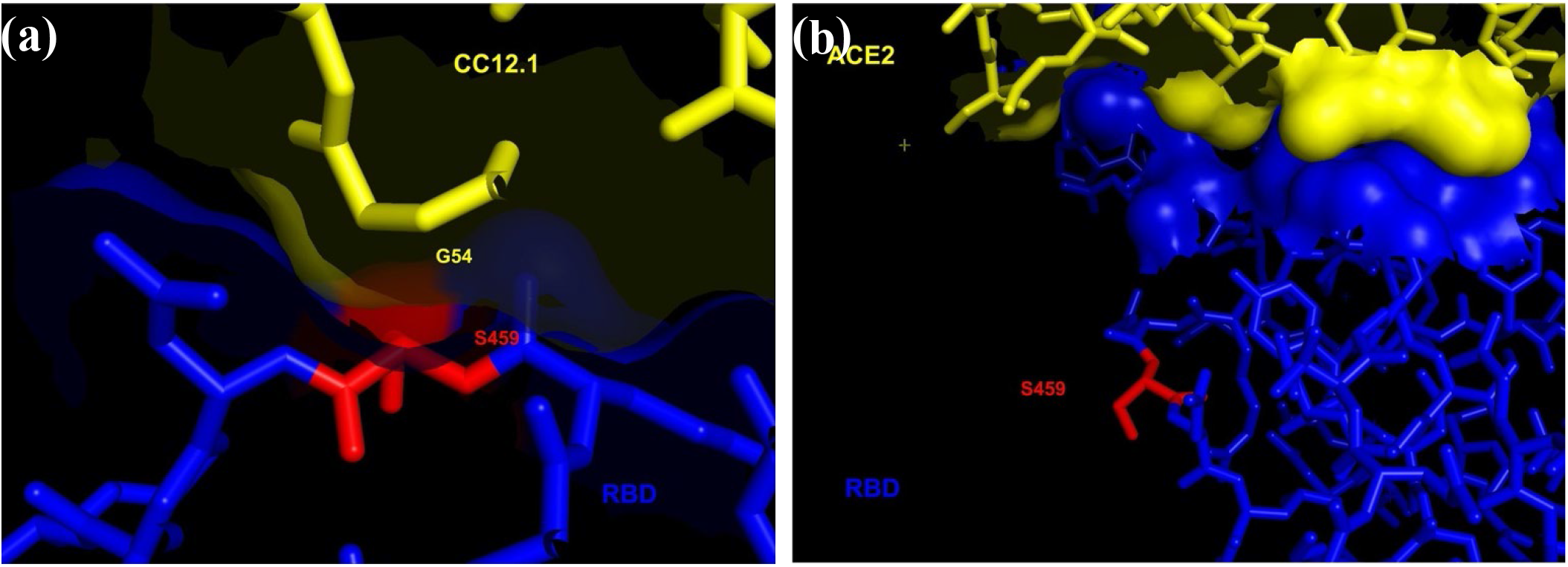
S459Y. (a) RBD:CC12.1 interface. (b) RBD:ACE2 interface.

A475 is in the interface with CC12.1 and on the edge of the interface with ACE2 (Fig 8). A475-O is a hydrogen bond acceptor for the hydrogens on T28-N and N32-Nδ2 in the heavy chain of CC12.1 (Fig 8a), and on S19-N in ACE2 (Fig 8b). A475V preserves the hydrogen bonding capability but significantly increases side chain volume in a region where packing is tight. There is little room for larger side chain around A475 in the interface with CC12.1, so this mutation is likely to be disruptive. A larger side chain appears much less likely to disrupt the interface with ACE2.

**Figure 8:**
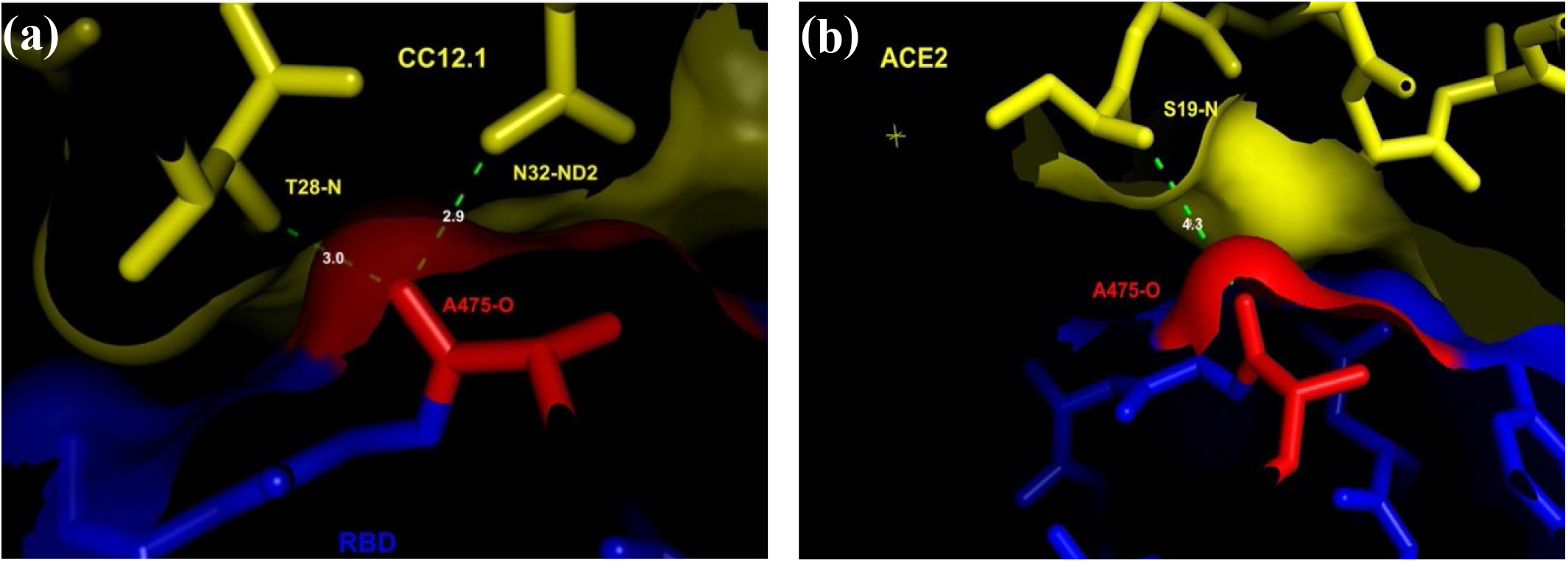
A475V. (a) RBD:CC12.1 interface. (b) RBD:ACE2 interface.

G476 is in the interface with CC12.1 and on the edge of the interface with ACE2 (Fig 9), making VdW contact with the aliphatic side chains of L27 in the heavy chain of CC12.1 (Fig 9a), and Q24 in ACE2 (Fig 9b). G476S is likely to disrupt the interface with CC12.1 due to its larger size and the introduction of unsatisfied hydrogen bond partners, but be permissive on the edge of the interface with ACE2.

**Figure 9:**
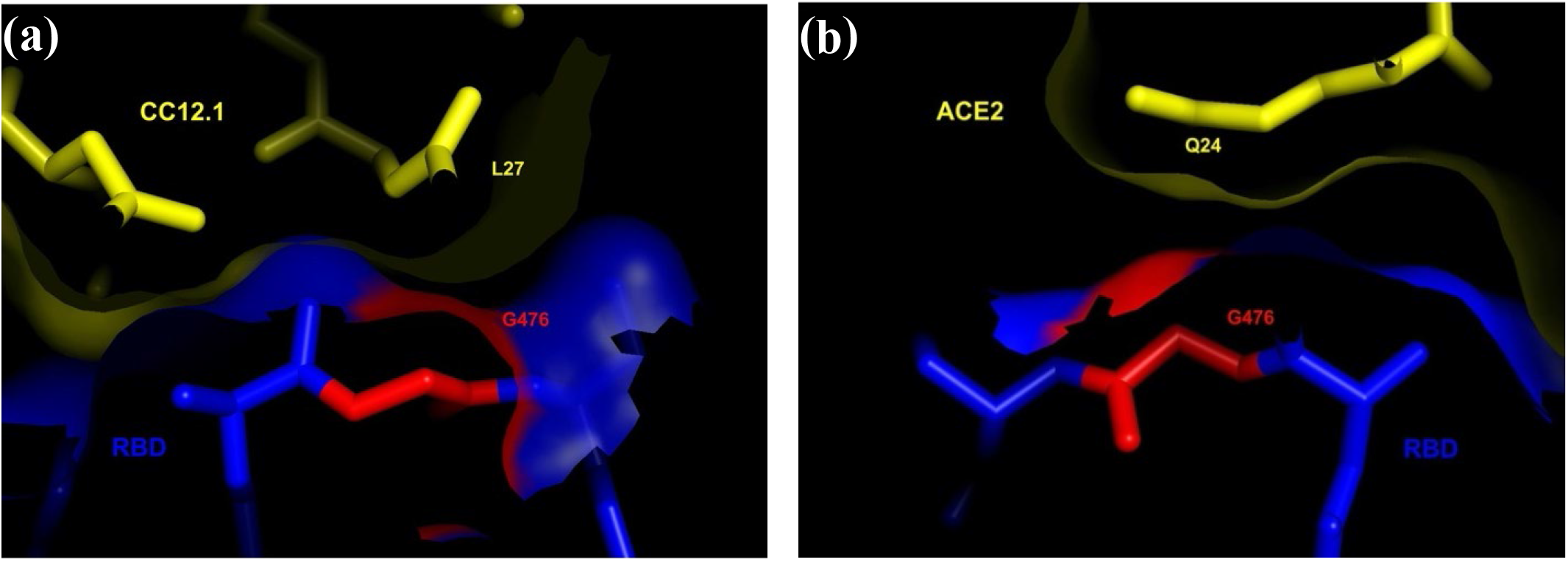
G476S. (a) RBD:CC12.1 interface. (b) RBD:ACE2 interface.

Q493 is in the interfaces with CC12.1 and ACE2 (Fig 10). Its side chain forms a hydrogen bond with Y99-Oη and makes VdW contact with the side chain of V98 across the interface with CC12.1 (Fig 10a). The Q493L mutation is approximately isosteric, but cannot form hydrogen bonds. Therefore, it is likely to be destabilizing to the interface with CC12.1. In the RBD:ACE2 interface, Q493-Nη2 exhibits static disorder, forming hydrogen bonds with either K31-Nζ or E35-Oζ1 (Fig 10b). However, K31-Nζ and E35-Oζ1 may form a salt bridge, as they do in crystal structure of the SARS-CoV spike protein.^20^ Therefore, Q493L is likely to be permissive in the interface with ACE2.

**Figure 10:**
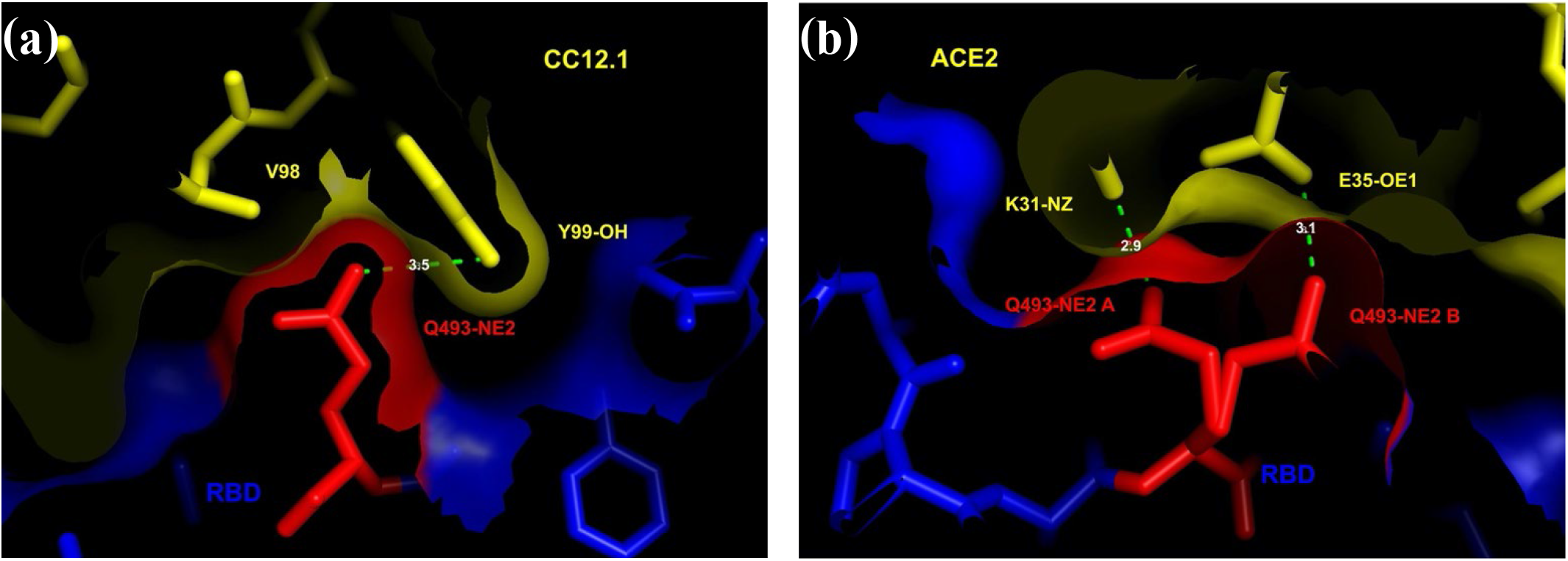
Q493L. (a) RBD:CC12.1 interface. (b) RBD:ACE2 interface.

N501 is in the interfaces with CC12.1 and ACE2 (Fig 11). N501-Nδ2 is hydrogen-bonded to G496-O in the RBD, while N501-Oδ1 forms a hydrogen bond with S30-Oγ in the CC12.1 light chain (Fig 11a). The N501Y mutation is likely to be disruptive in the interface with CC12.1 because it substitutes a much larger side chain that is unlikely to form similar hydrogen bonds. N501-Oδ1 forms a hydrogen bond with Y41-Oη, and interacts sterically with K353 in ACE2 (Fig 11b, c). N501Y may stabilize the interface with ACE2 by forming an edge-to-face interaction with Y41 and interacting sterically with K353 in ACE2. It should be noted that this type of steric interaction appears to be stabilizing in SARS-CoV binding to ACE2 because it supports a salt bridge formation between K353 and D38 in ACE2.^20^

**Figure 11:**
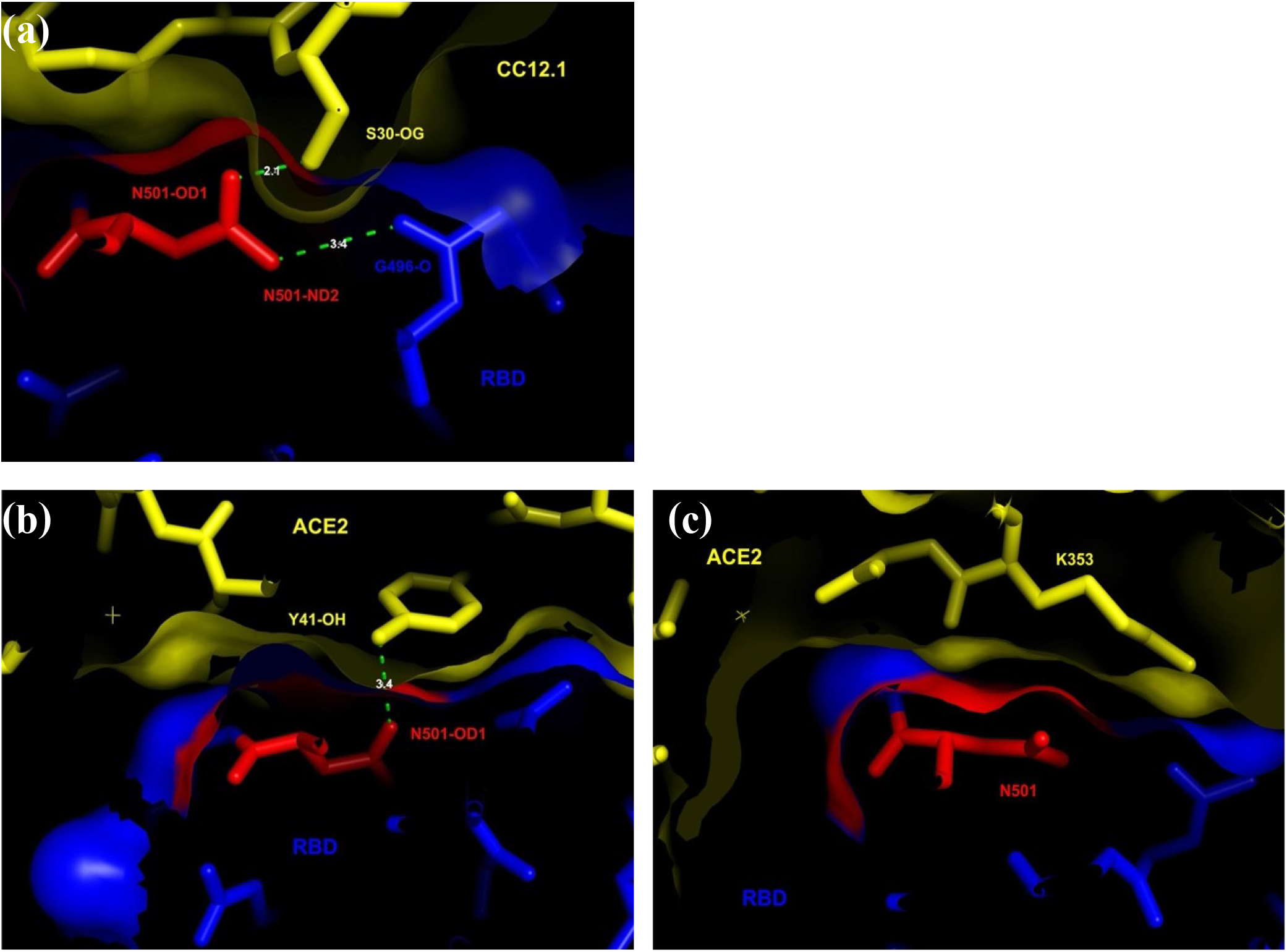
N501Y. (a) RBD:CC12.1 interface. (b) Hydrogen bond to Y41 In the RBD:ACE2 interface. (c) Interaction with K353 In the RBD:ACE2 interface.

Y505 is in the interfaces with CC12.1 and ACE2 (Fig 12). In the interface with CC12.1, Y505-Hη forms a hydrogen bond with the main chain of L91 and the sidechain of Q90 (Fig 12a). In the interface with ACE2, it forms a hydrogen bond with the sidechain of E37 (Fig 12b). A Y505C mutation can be accommodated sterically in this position but will not support these hydrogen bonds. L91 is not accessible to water, so Y505C is likely to be destabilizing in the interface with CC12.1. E37 is likely to be permissive on the edge of the interface with ACE2.

**Figure 12:**
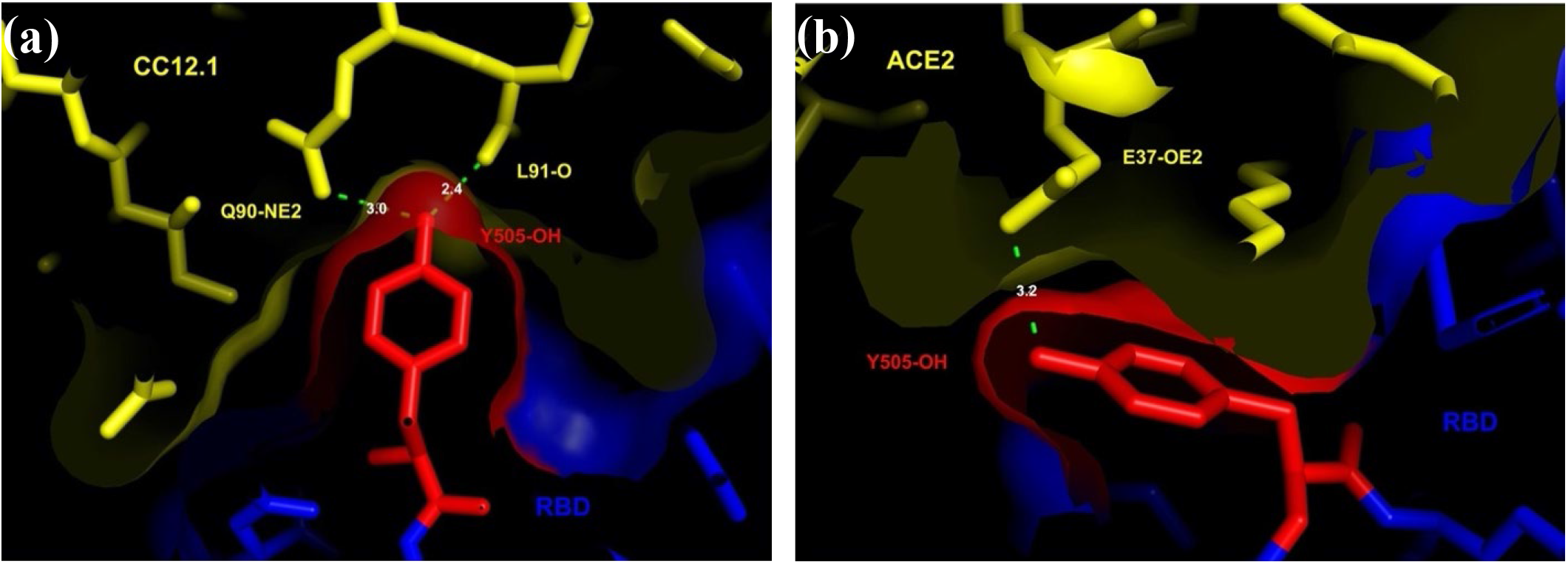
Y505C. (a) RBD:CC12.1 interface. (b) RBD:ACE2 interface.

### Hotspots on the edge of the RBD:CC12.1 interface (table 2)

S477, E484, F490, S494, and V503 are all on the edge of the interface with CC12.1 (Figs. 13-17). None of them are in the interface with ACE2, although E484, F490, and V503 are on the edge of the interface with ACE2. The number of mutations in these residues among infectious viruses suggests that mutations are readily tolerated on the edges of the interface with ACE2. Upon visual inspection, it is not clear that any of these mutations would disrupt the interface with CC12.1, and the permissive nature of these sites for ACE2 binding suggests that they are all likely to be permissive for CC12.1 binding.

### Hotspots in the RBD:P2B-2F6 interface (table 3)

G446 is in the interfaces with P2B-2F6 and ACE2 (Fig 18). The G446 main chain makes VdW contact with the Y27 side chain the P2B-2F6 heavy chain (Fig 18a), and both G446V and G446S would likely disrupt this interface. There is more room for a side chain in the interface with ACE2, making both mutations permissive.

**Figure 13:**
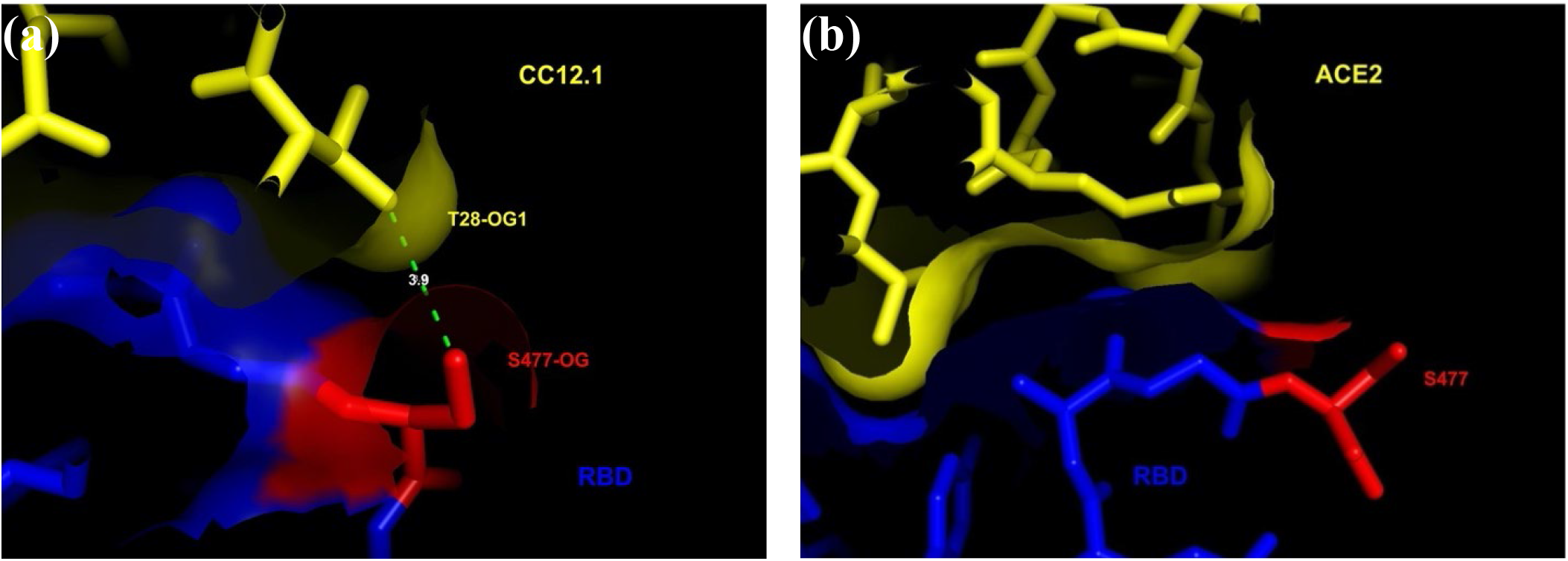
S477N/G/R/I. (a) RBD:CC12.1 interface. (b) RBD:ACE2 interface.

**Figure 14:**
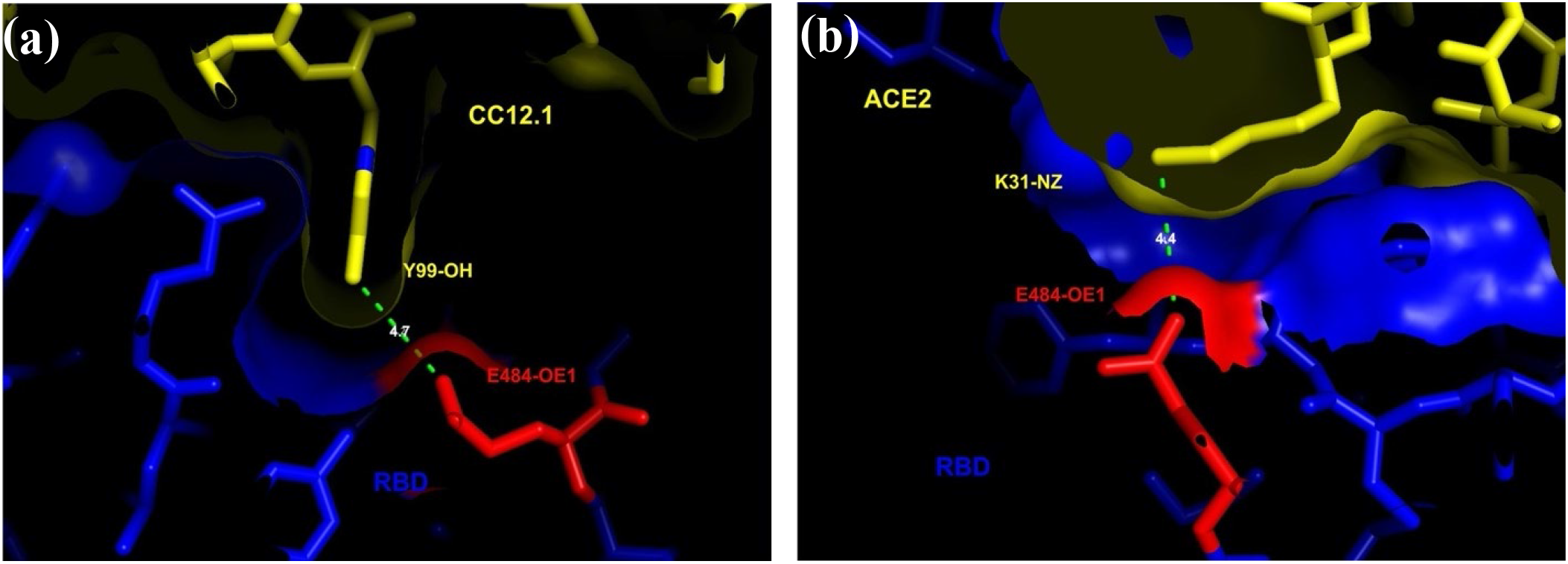
E484Q/K/A. (a) RBD:CC12.1 interface. (b) RBD:ACE2 interface.

**Figure 15:**
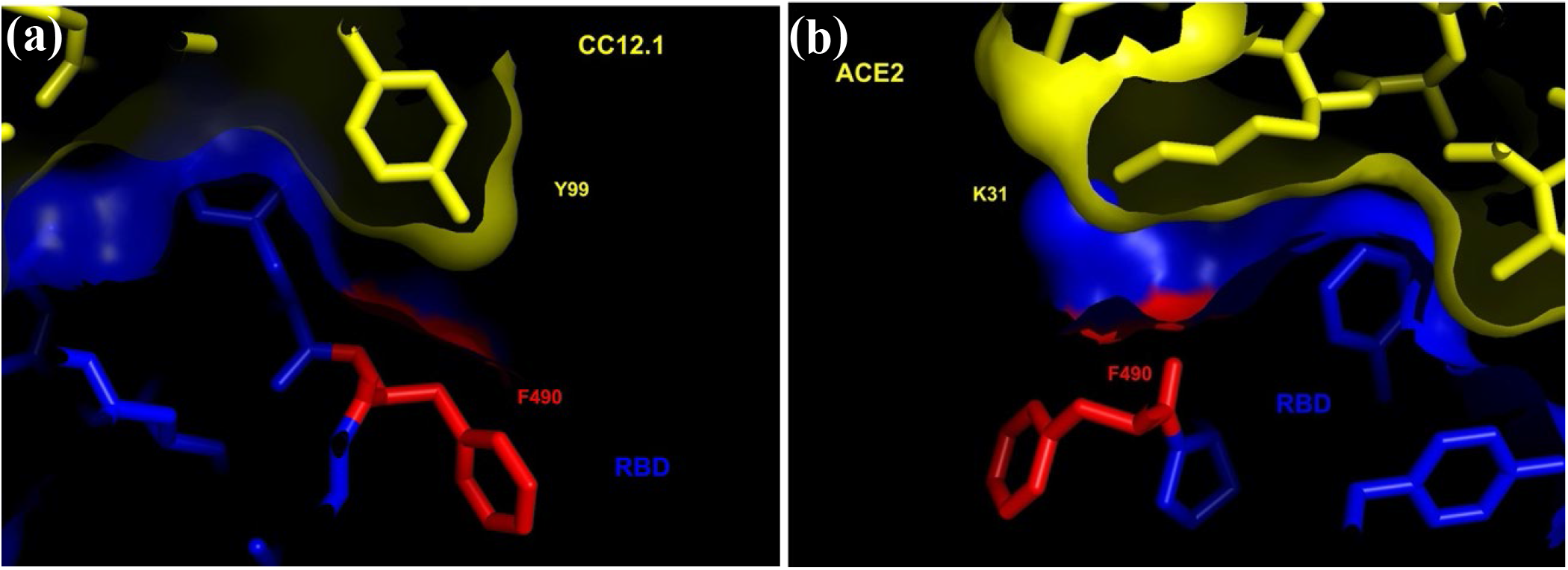
F490S/L. (a) RBD:CC12.1 interface. (b) RBD:ACE2 interface.

**Figure 16:**
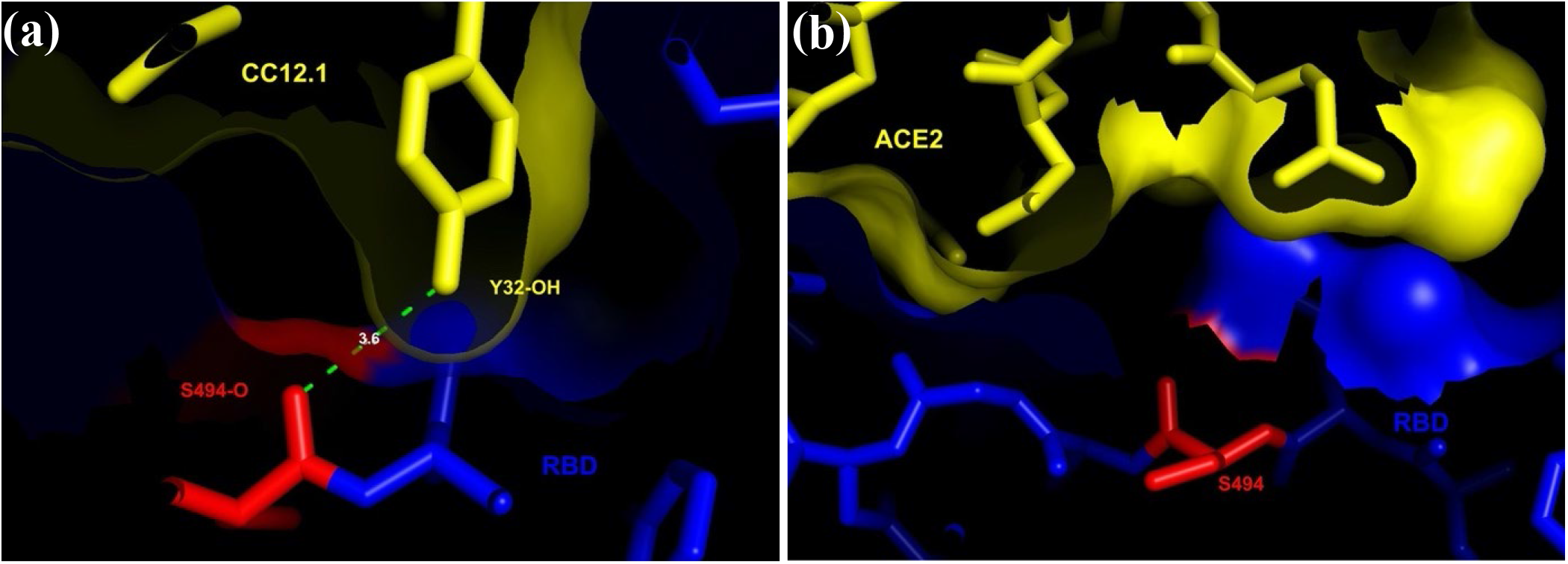
S494P/L. (a) RBD:CC12.1 interface. (b) RBD:ACE2 interface.

**Figure 17:**
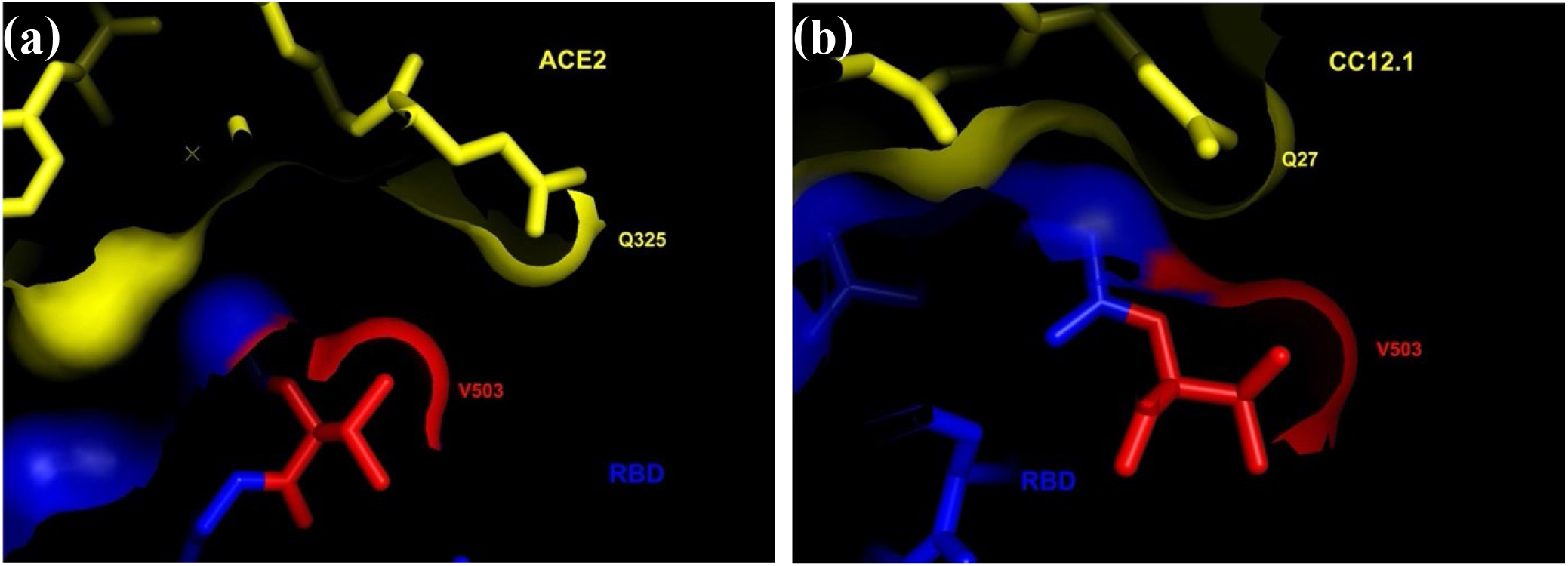
V503F. (a) RBD:CC12.1 interface. (b) RBD:ACE2 interface.

**Figure 18:**
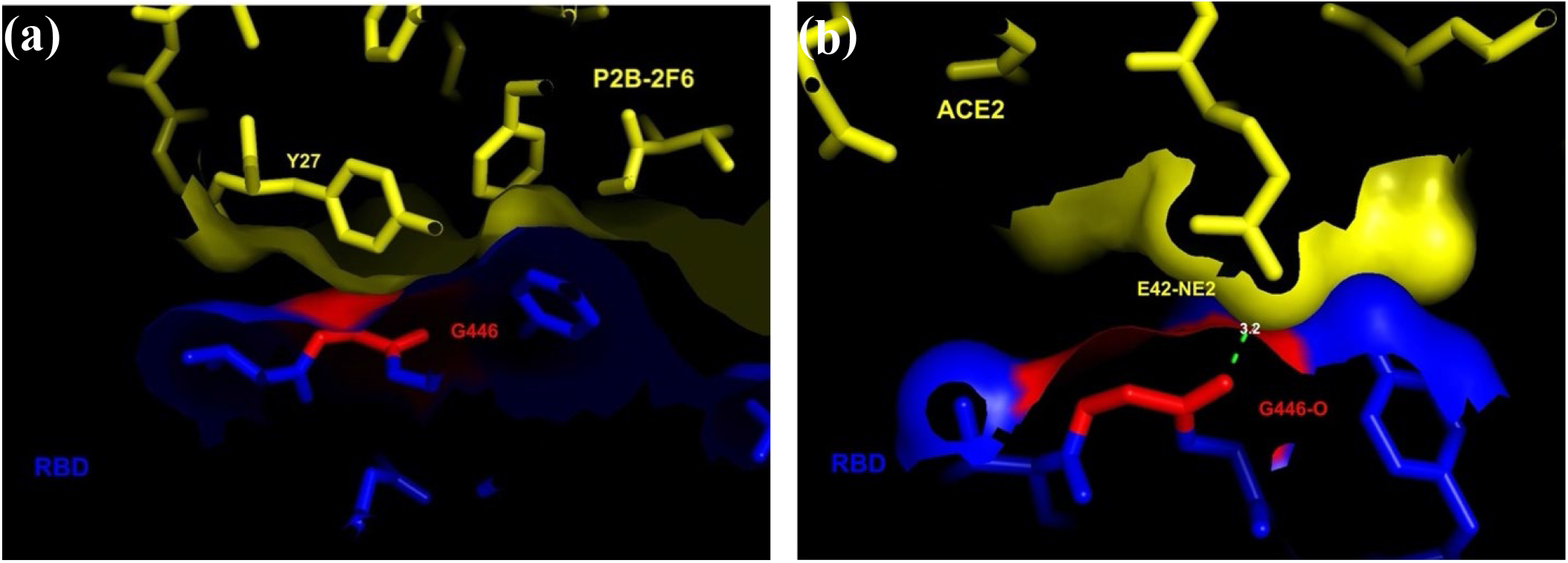
G446V/S. (a) RBD:P2B-2F6 interface. (b) RBD:ACE2 interface.

L452 is in the interface with P2B-2F6 but not ACE2 (Fig 19). The sidechain of L452 makes VdW contact with the side chains of I103 and V105 in the P2B-2F6 heavy chain (Fig 19a), and L452M is likely to be permissive.

**Figure 19:**
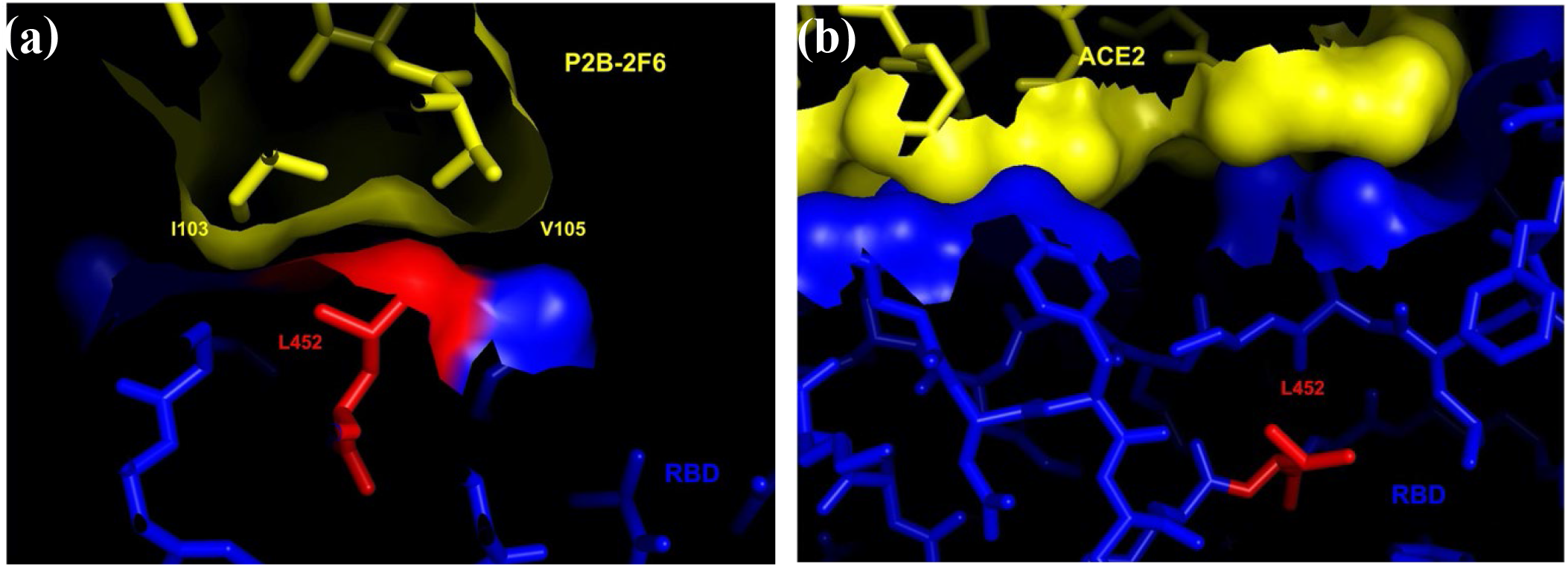
L452M. (a) RBD:P2B-2F6 interface. (b) RBD:ACE2 interface.

V483 is in the interface with P2B-2F6 but not ACE2 (Fig 20). The sidechain of V483 makes VdW contact with the G31 main chain in the P2B-2F6 light chain (Fig 20a). It is likely that V483A and V483I may be accommodated at this position and permissive to the interface with P2B-2F6, but V483F is likely to disrupt the interface with P2B-2F6 due to its greater side chain volume.

**Figure 20:**
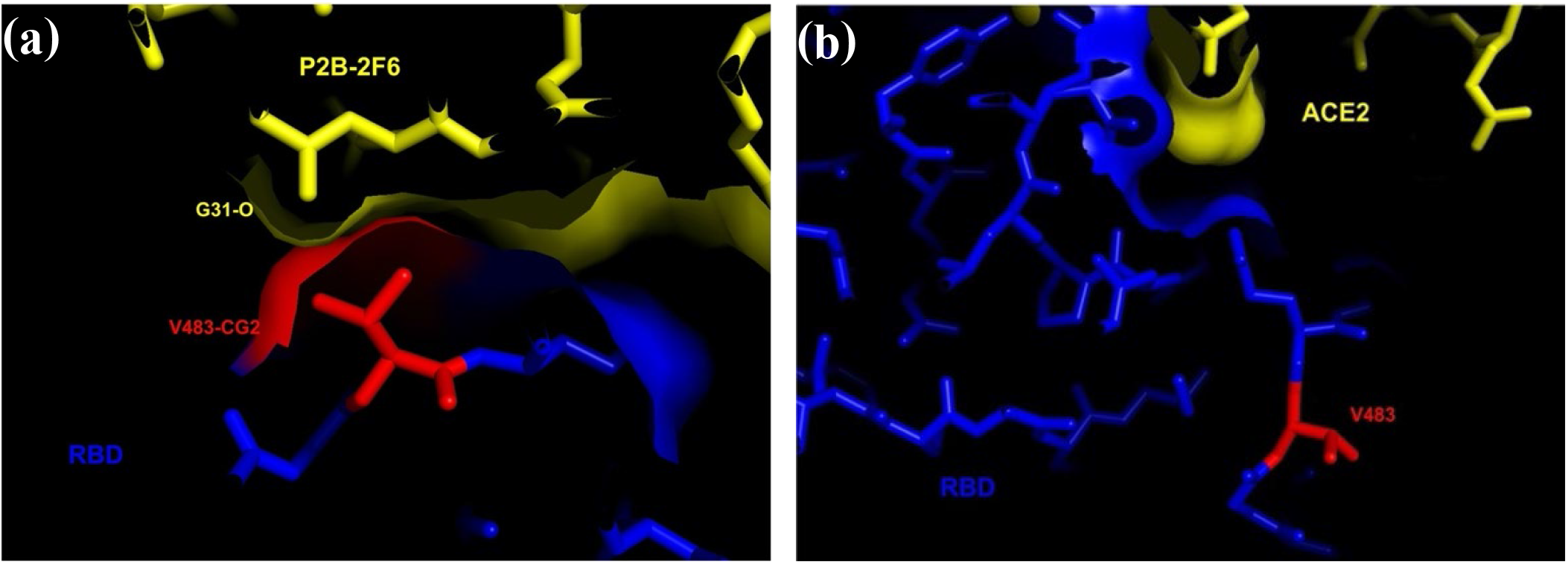
V483A/F/I. (a) RBD:P2B-2F6 interface. (b) RBD:ACE2 interface.

E484 is in the interface with P2B-2F6, and on the edge of the interface with ACE2 (Fig 21). E484-OE2 forms hydrogen bonds with Y34-Oη of the P2B-2F6 light chain, and with R112-Nη1 of the P2B-2F6 heavy chain (Fig 21a). E484-O forms a hydrogen bond with N33-Nδ2 of the P2B-2F6 light chain (Fig 21b). In the interface with ACE2, E484 and K31 form a salt bridge (Fig 21c). These interactions suggest that E484A, E484Q, and E484K will all be destabilizing in the interface with P2B-2F6, but permissive in the interface with ACE2.

**Figure 21:**
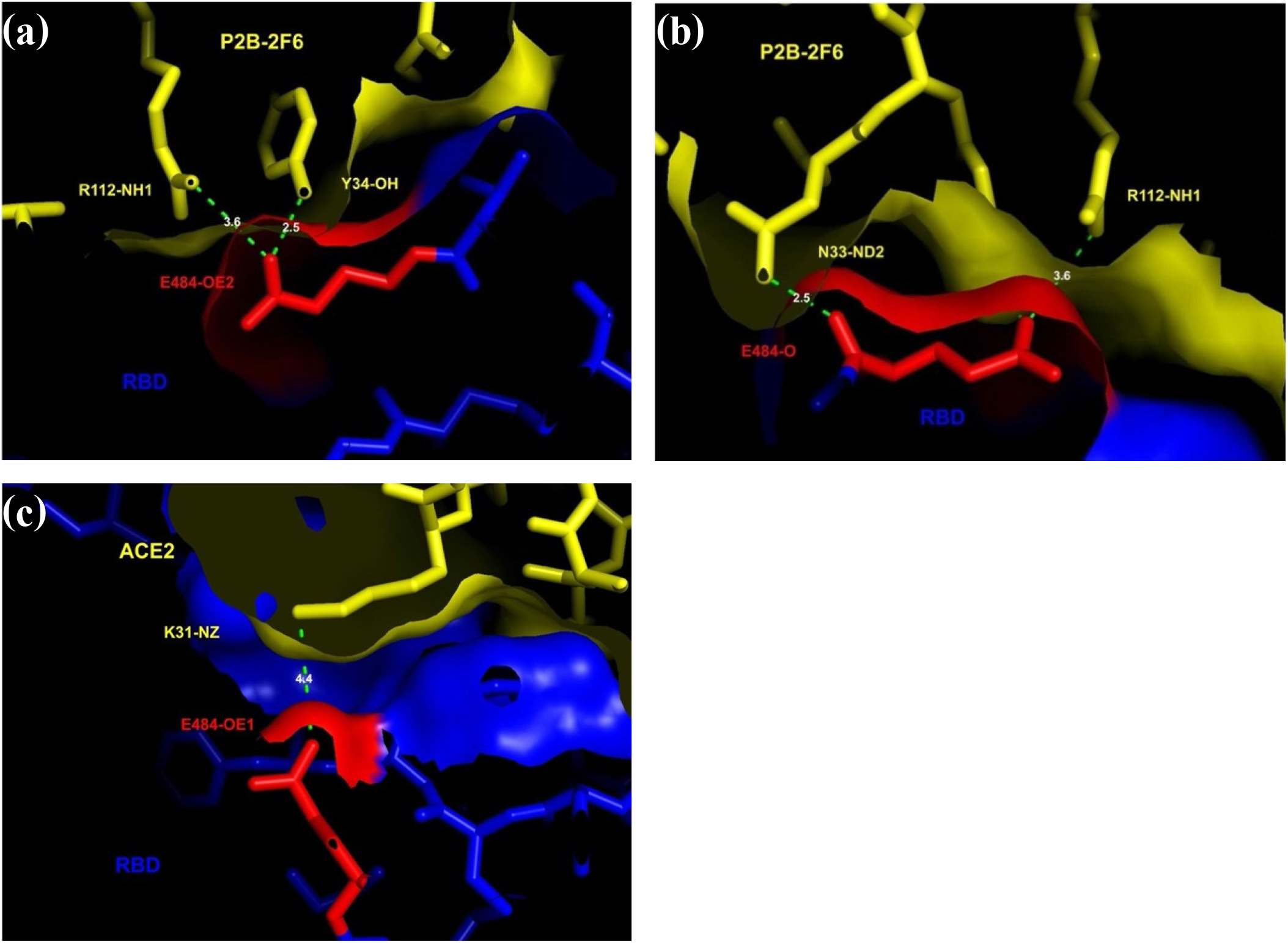
E484Q/K/A (a) Hydrogen bond to R112 and Y34 In the RBD:P2B-2F6 interface. (b) Hydrogen bond to N33 and R112 In the RBD:P2B-2F6 interface. (c) RBD:ACE2 interface.

F490 is in the interface with P2B-2F6, but not the interface with ACE2 (Fig 22). The F490 sidechain makes VdW contact with P107 and V105 in the P2B-2F6 heavy chain (Fig 22a), and the K31 sidechain in ACE2 (Fig 22b). F490S can be accommodated sterically, but leaves hydrogen bond partners buried and unsatisfied. F490L can also be accommodated sterically, without these destabilizing features. Thus, both mutations will likely be permissive in the interface with P2B-2F6.

**Figure 22:**
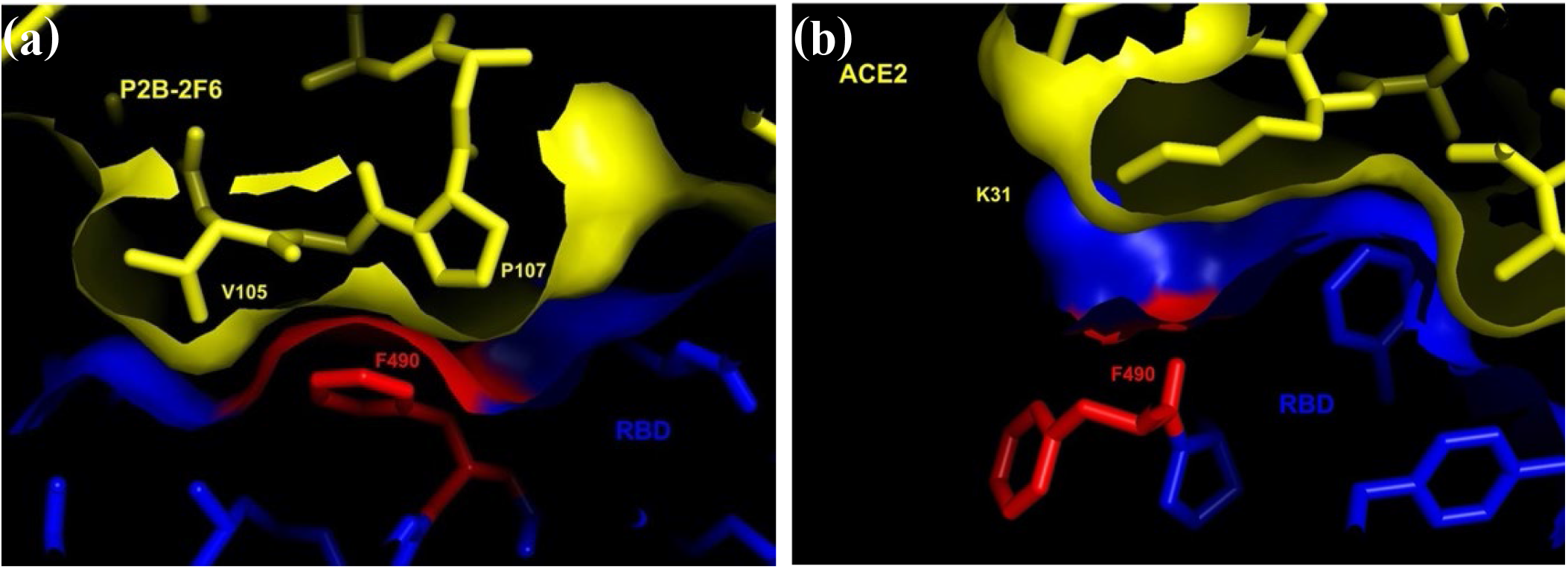
F490S/L. (a) RBD:P2B-2F6 interface. (b) RBD:ACE2 interface.

### Hotspots on the edge of the RBD:P2B-2F6 interface (table 4)

R346K, K444, V445A, G482S, G485R, Q493L, and S494P/L are all on the edge of the interface with P2B-2F6 where they are likely to be quite tolerant of mutations (Figs. 23-29). Only Q493 is in the interface with ACE2 and, for reasons given above, Q493L is likely to be permissive in this interface.

**Figure 23:**
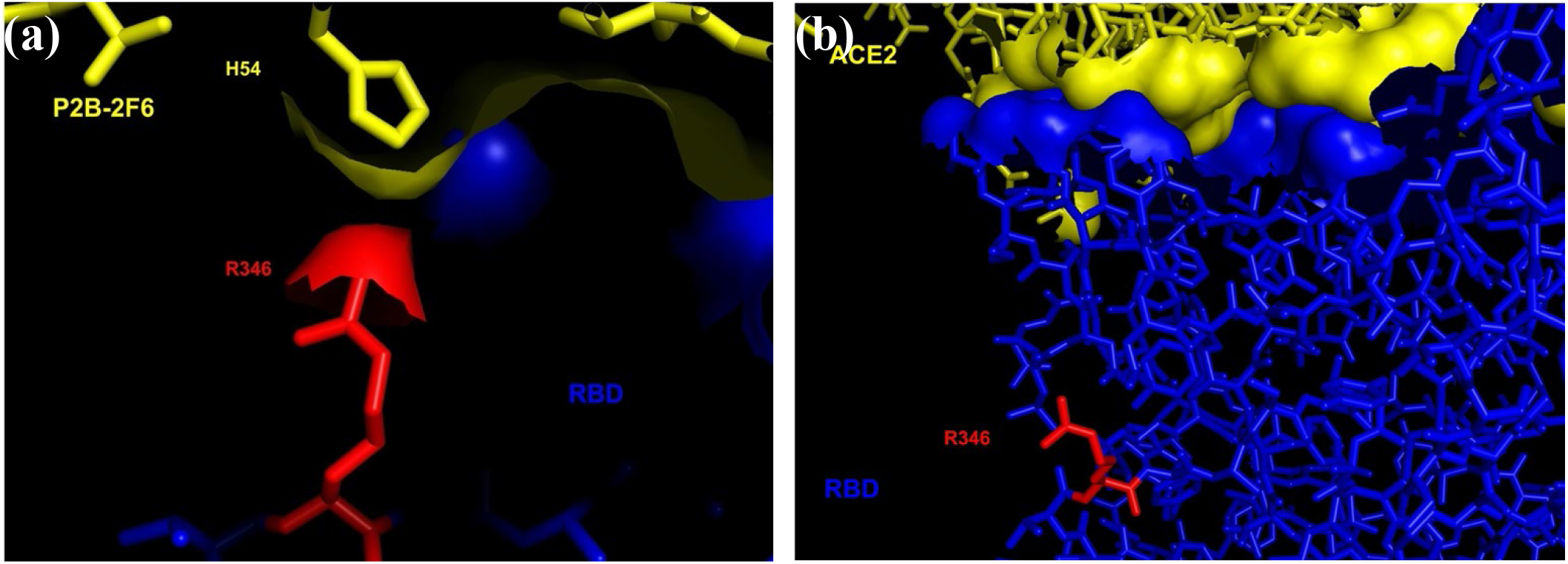
R346K. (a) RBD:P2B-2F6 interface. (b) RBD:ACE2 interface.

**Figure 24:**
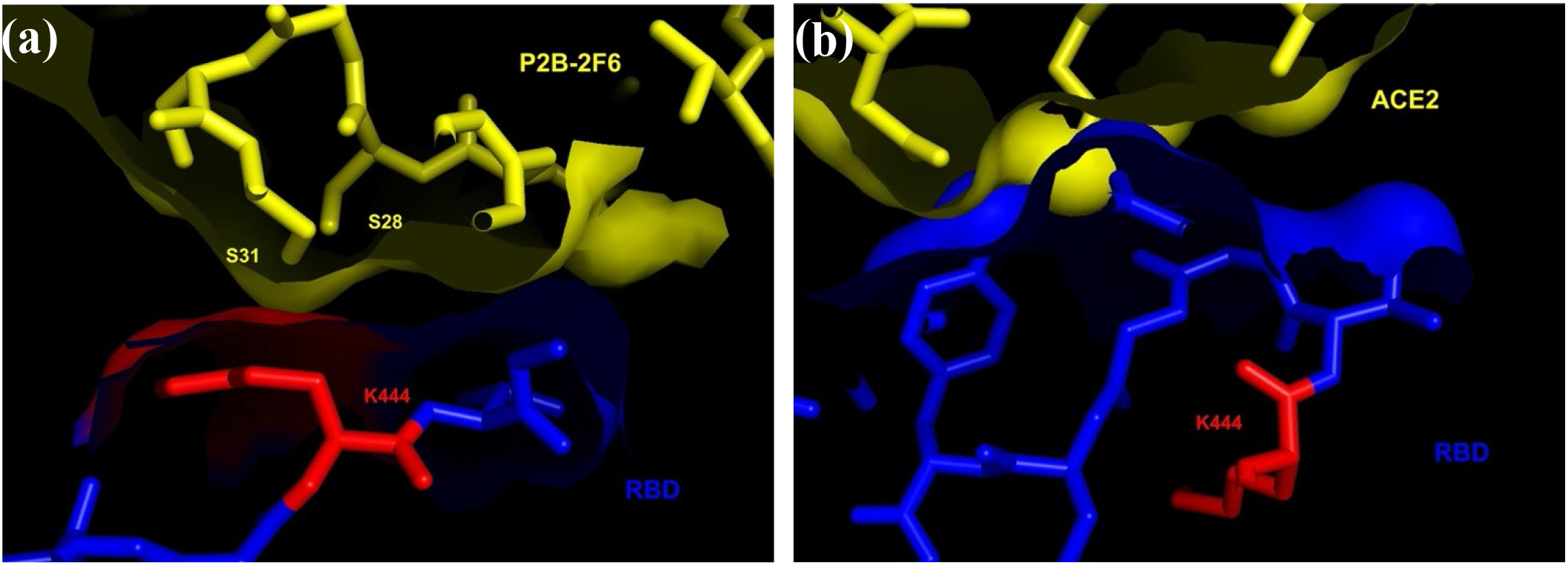
K444N/R. (a) RBD:P2B-2F6 interface. (b) RBD:ACE2 interface.

**Figure 25:**
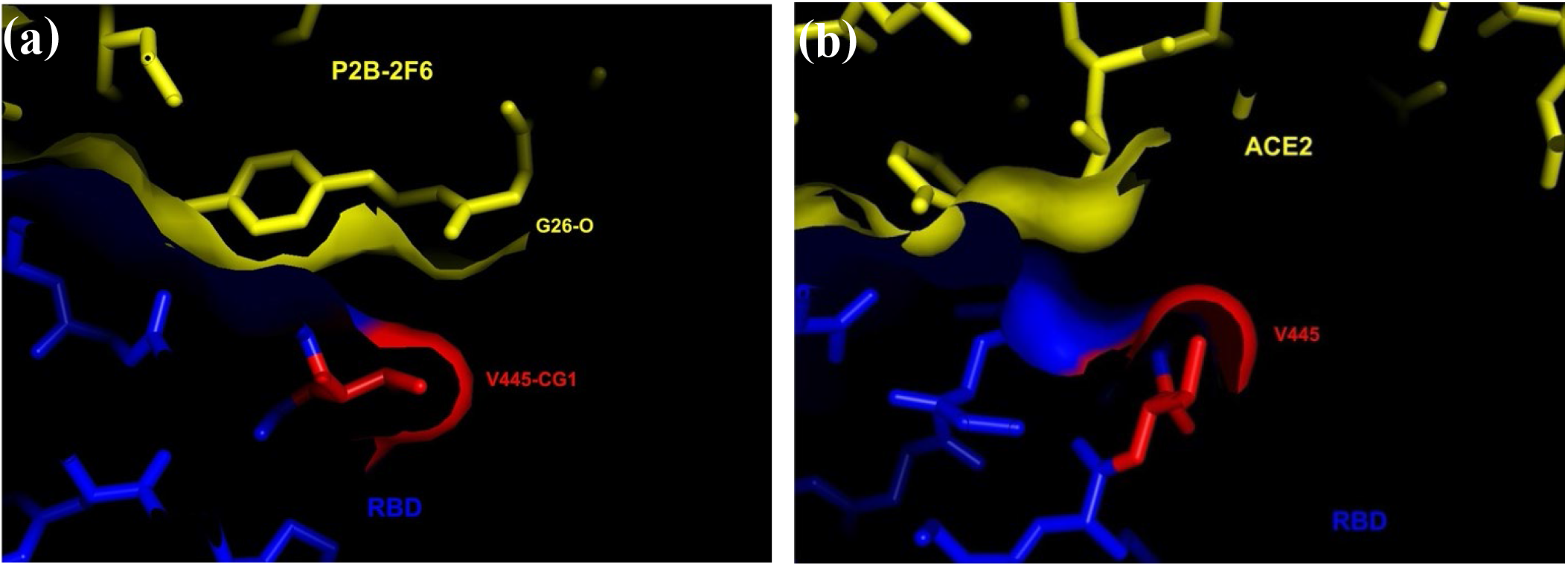
V445A. (a) RBD:P2B-2F6 interface. (b) RBD:ACE2 interface.

**Figure 26:**
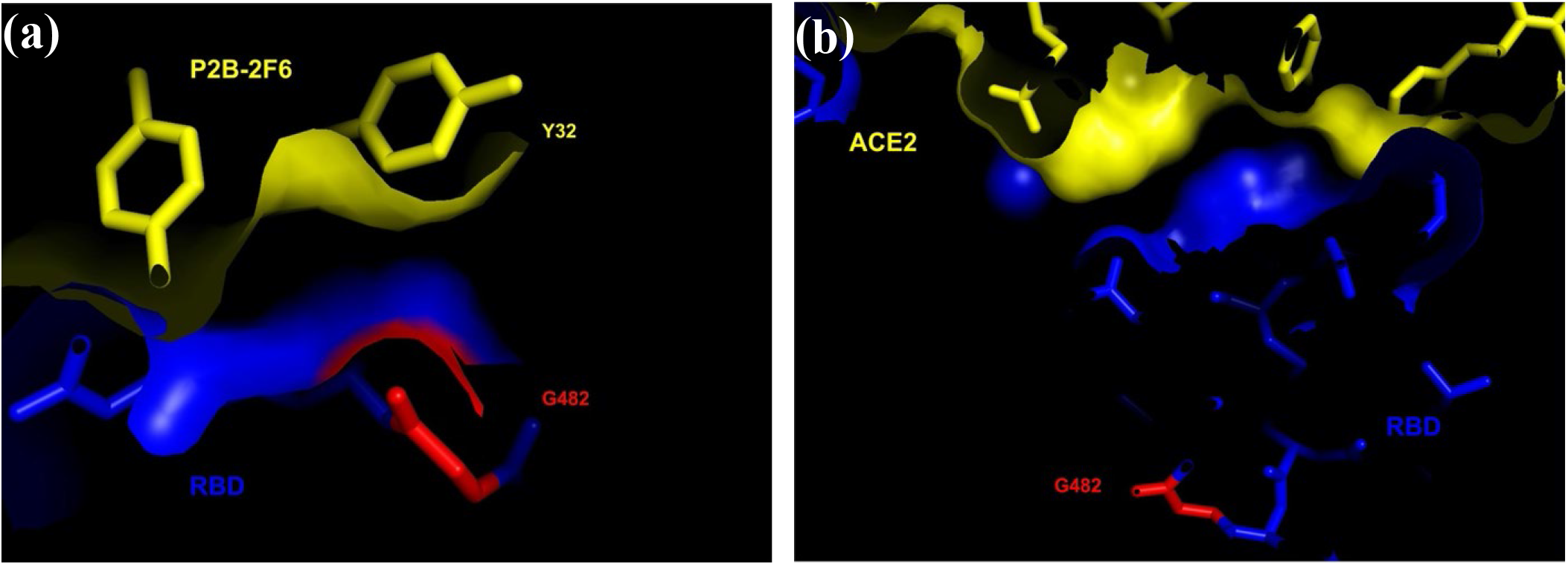
G482S. (a) RBD:P2B-2F6 interface. (b) RBD:ACE2 interface.

**Figure 27:**
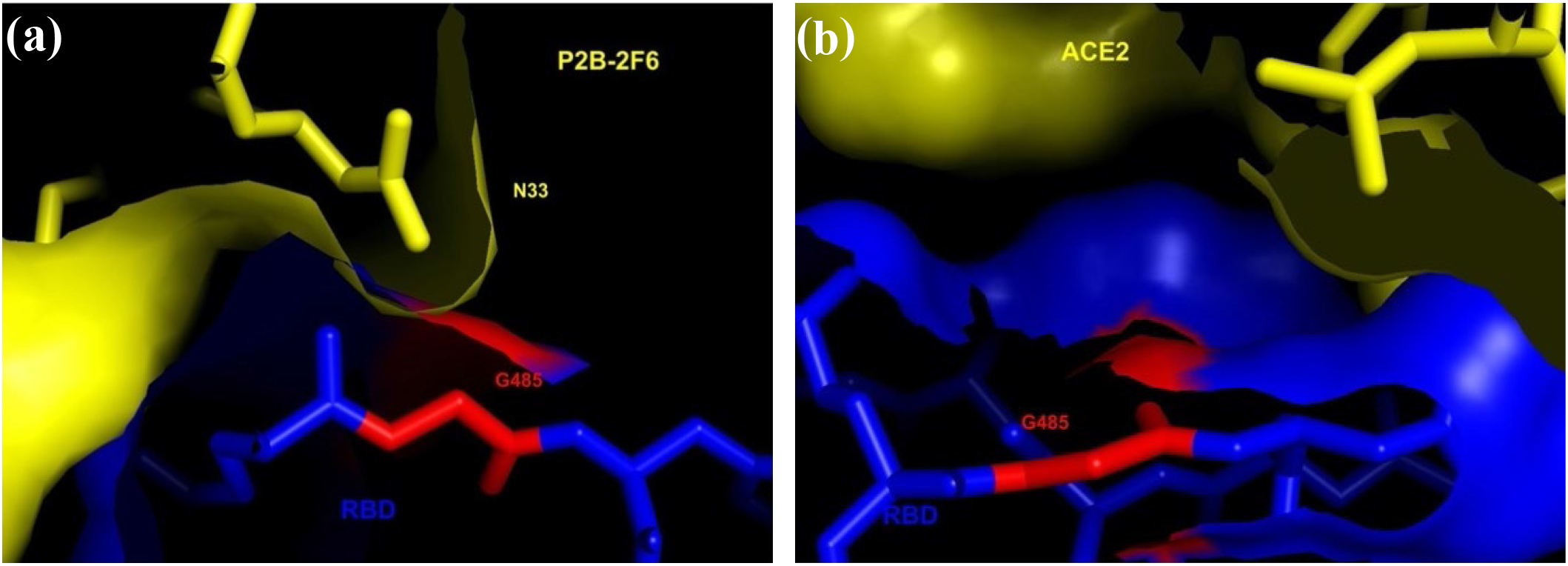
G485R. (a) RBD:P2B-2F6 interface. (b) RBD:ACE2 interface.

**Figure 28:**
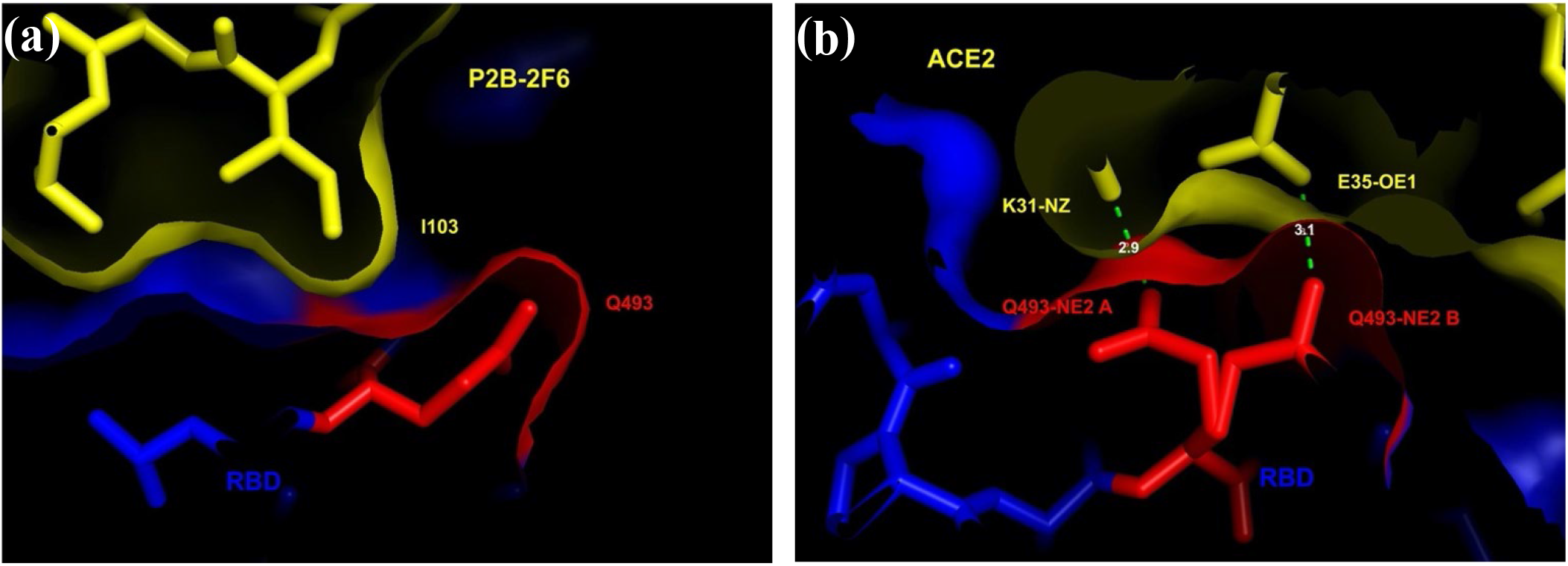
Q493L. (a) RBD:P2B-2F6 interface. (b) RBD:ACE2 interface.

**Figure 29:**
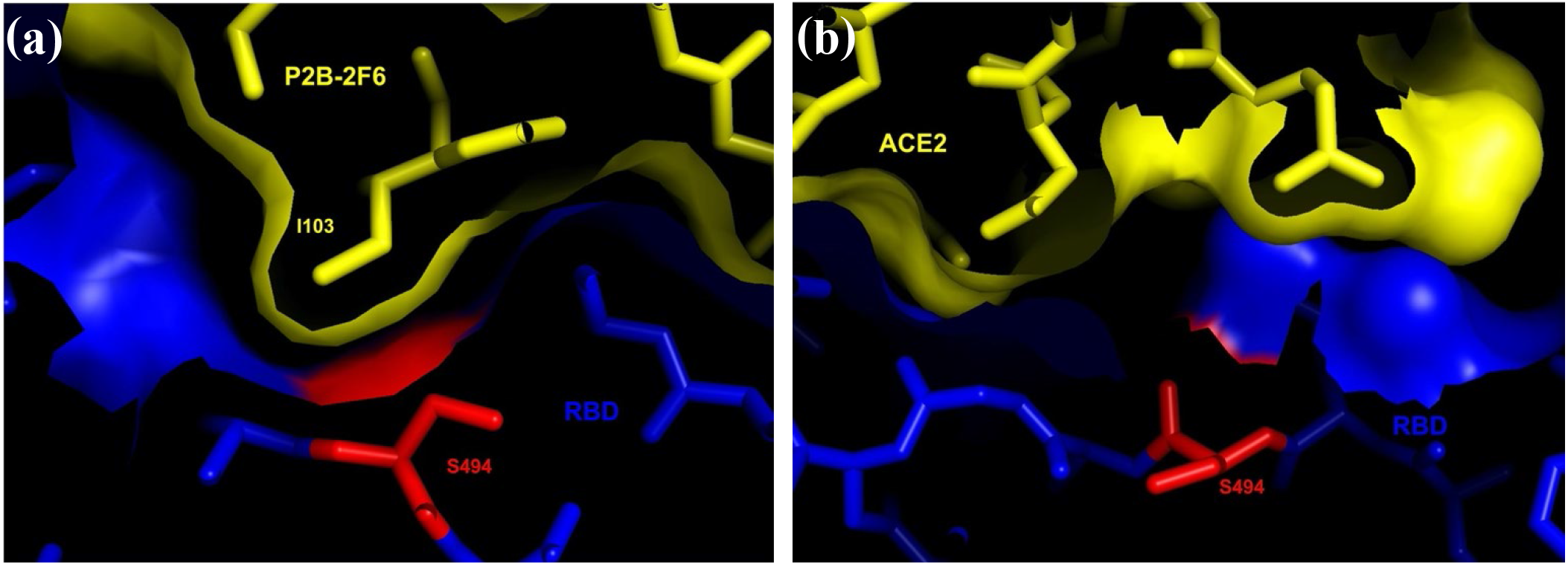
S494P/L. (a) RBD:P2B-2F6 interface. (b) RBD:ACE2 interface.

## Discussion

The spike protein mutations listed in tables 1-4 represent infectious SARS-CoV-2 variants in which the replicative and pathogenic mechanisms of the virus are functional. The first step of SARS-CoV-2 infection is the attachment of its spike protein to ACE2. Therefore, it is not surprising that the mutations examined in this study either did not involve portions of the spike protein that interface with ACE2, or they were structurally or functionally conservative so as to not disrupt the RBD:ACE2 interface.

Functionally important regions of the spike protein are ideal targets for neutralizing antibodies because they tend to be highly conserved, and intolerant of mutations that would interfere with immune recognition by antibodies. The two antibodies chosen for study were of interest because high resolution structures of their binding interfaces with the spike protein were available, and because their binding interfaces overlapped with the RBD:ACE2 binding interface. Unfortunately, both the RBD:CC12.1 and the RBD:P2B-2F6 binding interfaces also involve residues outside the RBD:ACE2 interface, and most of the mutations listed in tables 1-4 are likely to destabilize or disrupt the RBD:antibody interfaces in such a way that ACE2 binding is preserved.^21^

There are two important implications to this pattern. First, it suggests that mutated forms of SARS-CoV-2 may be able to reinfect people who have recovered from an earlier infection. Natural infection will tend to generate multiple antibodies to distinct aspects of the spike protein, and immune evasion by mutation becomes increasing unlikely as the number of independent neutralizing antibodies increases, especially since SARS-CoV-2 has a relatively low genetic diversity.^22^ However, nearly all of the mutations listed in tables 1-4 were identified multiple times on different continents, presumably in the absence of evolutionary pressure, suggesting that they represent “hotspots” where mutation is spontaneous and frequent. It should be noted that all but one of the mutations involved a single nucleotide substitution (R403M involved two nucleotides). When evolutionary pressure is applied, we must expect the potential for immune evasion to increase.

The second implication of this pattern is that vaccines containing only one variant of the spike protein as an immunogen may induce little or no protection against variant strains that are already circulating among highly mobile human populations. An inability to protect against variant strains may severely limit the strategic benefit of vaccines containing only the original strain. To have long-term efficacy against SARS-CoV-2 it may be necessary for vaccines to include multiple variants of the spike protein as immunogens.^23,24^ This strategy is routinely applied against influenza, and all formulations in recent years have all included three or four variant strains. In the case of influenza, however, it has been possible to predict which strains will become epidemic, and the antigenic features of those strains tend to be stable for a season. SARS-CoV-2 differs in that many variant strains are already circulating, and their antigenic features are not stable.

These implications are supported by experimental results recently described in preprint form that examined the ability of artificially-generated single site mutations to escape recognition by a panel of 10 antibodies not considered above.^25^ In some cases, the mutations considered in that work were the same as the naturally occurring mutations considered above, and they were shown to reduce the ability of an antibody to bind ACE2 and to neutralize the variant virus.

A limitation of this study is that the ability of a mutation to disrupt, destabilize, or permit a protein-protein interface was gauged visually, without the benefit of a molecular mechanics force field to quantify the effects of mutation, or of experimental data to verify the expected effects. Consequently, individual conclusions about a specific mutation may be in error. However, the overall pattern – namely that mutations at the antibody interfaces may not be permissive while not interfering with the ACE2 interface – is sufficiently clear and strong that sporadic errors would not negate the principle conclusions.

